# EffectorGeneP: accurate gene annotation in pathogen genomes from infection transcriptomes

**DOI:** 10.64898/2026.05.04.722827

**Authors:** Jana Sperschneider, Camilla Langlands-Perry, Jian Chen, Jibril Lubega, Taj Arndell, David C. Lewis, Eva C. Henningsen, Cheryl Blundell, Thomas Vanhercke, Kostya Kanyuka, Melania Figueroa, Peter N. Dodds

## Abstract

Accurate gene annotation is crucial for inference of biological knowledge from genomes. However, non-canonical genes such as orphan or single-exon genes as well as those residing in rapidly evolving regions are routinely dismissed in annotation pipelines. In filamentous pathogen genomes, this disproportionately affects the annotation of genes encoding disease-promoting effector proteins. We introduce EffectorGeneP, a machine learning tool that self-trains on transcript data, predicts the most likely coding sequence from transcripts and effectively separates *bona fide* genes from transcriptional noise. EffectorGeneP annotates over 95% of known effectors correctly, while other state-of-the-art methods annotate 15%-78%. We show that EffectorGeneP expands the predicted secretome of pathogens by over 50% and that high-throughput screening of an effector library in plant protoplasts uncovers the previously poorly annotated *AvrSr26* gene family in the wheat stem rust fungus. EffectorGeneP decodes genomes at unprecedented resolution and will enable the study of biological processes in important pathogen species.

## Introduction

Understanding genome content and deciphering the role of functional elements relies on accurately locating and designating genes within DNA sequences (gene annotation). Gene annotation is typically achieved either via *ab initio* methods or through approaches that utilize homology-based searches or transcriptional data. Whilst transcriptome-based methods consistently outperform *ab initio* methods^1^, gene annotation is still erroneous and at its core relies on decades-old approaches such as Hidden Markov Models (HMMs) that in most cases do not fully utilize transcriptional data. Recently, *ab initio* machine learning methods that incorporate a HMM layer have been released, Helixer^2,3^ and Tiberius^4^. However, a limited number of species are currently in the training sets, re-training deep learning models is GPU-dependent and a high-quality reference annotation is required for training.

Apart from computational challenges, gene annotation is complicated by heterogeneity in the gene pool of a species. Genes encode proteins of various sizes and amino acid composition, can reside in genomic regions with different properties, can deviate from canonical regulatory sequence motifs or splice sites and can be either conserved or unique to a species or an isolate (orphan genes). Furthermore, genomes contain numerous non-functional small ORFs, while long ORFs are unlikely to occur by chance. Thus, both length filters and homology-based evidence are heavily used by most annotation methods. As a consequence of these constraints, annotation of non-canonical genes remains challenging^5^. Most pipelines filter for highly conserved genes to use as their training set, leading to a bias against the signatures of orphan genes^6,7^. Similarly, small genes and single-exon genes are often removed from training sets or gene predictions to minimize false positive annotations. For example, CodingQuarry only allows single-exon genes larger than 600 nucleotides (nts) for training^8^ and Tiberius removes genes with a coding length of less than 200 nts from its final set of predictions^4^. For the annotation of non-canonical genes, transcriptional data can provide evidence for their existence as well as accurate information on intron-exon boundaries and alternative splicing.

The biological importance of non-canonical genes from both host and pathogen is increasingly recognised^9,10^. For example, pathogen effector genes encode secreted proteins that facilitate host colonization. Pathogen effectors have a broad functional range and are enriched for small proteins and orphan genes^11^. CodingQuarry is thus far the only annotation tool that has a specialized ‘pathogen mode’ (PM) for effector annotation^12^, which uses a GHMM framework with additional states to model the presence of a signal peptide and a codon model derived from genes encoding proteins with high-cysteine content and atypical codon usage. To improve the annotations of non-canonical genes and overcome the limitations of GHMMs, custom pipelines typically based on sequence or motif searches have been developed. For example, RxLR^13^ and Crinkler (CRN)^14,15^ effector genes are typically annotated from genomes using motif and signal peptide searches in open reading frames followed by manual inspection and curation which has led to hundreds of additional effector gene models in these species^15–17^. However, such methods are time-consuming, error-prone and restricted to certain pathogen species. To improve annotation of fungal effectors, assembled transcripts that do not contain annotated genes have been added to an existing annotation if the longest ORF encodes a protein with a signal peptide^18,19^. Whilst such methods can be fully automated, they do not include models for coding probabilities and might thus lead to false positive gene annotations. Furthermore, they do not correct a mis-annotated gene in the existing annotation. To resolve these difficulties, here we have developed a machine learning (ML) based approach called EffectorGeneP to enhance gene annotation and specifically the annotation of pathogen effector genes.

## Results

### Training of EffectorGeneP for classifying genomic features

We compiled a set of ten fungal species (Supplementary Table 1) that were chosen for having high-quality genome assemblies, *in planta* RNA-seq data and known effector genes. As a control, a non-pathogenic yeast species (*Saccharomyces cerevisiae*) was also included with RNA-seq data from *in vitro* culture. In gene annotation, it is essential to assess the coding potential of ORFs to distinguish those encoding actual proteins from non-coding ORFs which randomly occur in genomes. Small ORFs occur at high frequency in genomes with most of these encoding very small proteins with high variability in GC content (Figure 1). This is the reasoning behind the practice of most gene annotation tools to only annotate small single-exon genes if they have homology evidence, which can exclude genes encoding highly diverse pathogen effectors. To develop an alternative approach that is agnostic with respect to gene conservation, we used assembled transcripts from infection RNA-seq data to train ML models for each species (Supplementary Table 1). These ML models distinguish between the following types of genomic regions: coding sequences (CDS) of non-secreted proteins, CDS of secreted proteins, CDS of secreted effector proteins, 3’ and 5’ UTRs, intergenic sequences, introns, small intergenic ORFs, random CDS sequences and Kozak sequences. No existing gene annotation or homology filtering is required for generating the training sets. Classification accuracy of these models on several sequence sets demonstrated high prediction accuracy (Supplementary Table 2). EffectorGeneP classified ∼99% of the small ORFs and random CDS sequences correctly and for the reference annotation CDSs, ∼98% were classified as such, with the majority (70-80%) placed in the non-secreted protein class. Next, we demonstrate how to use the ML models to accurately annotate genes from transcriptional data alone.

**Figure 1:**
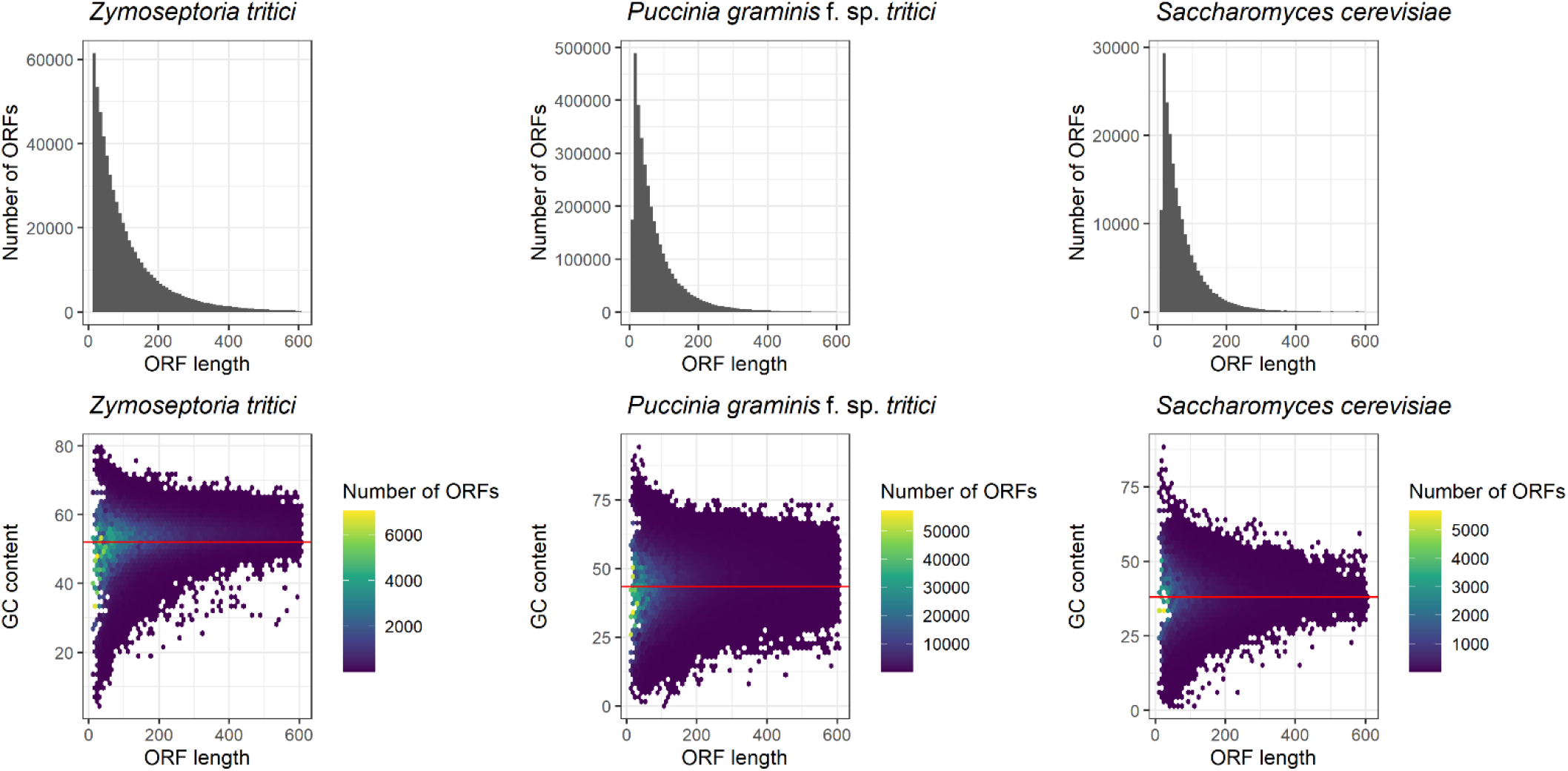
Small, complete ORFs occur at high frequency in fungal genomes. Three genomes are shown: *Zymoseptoria tritici* (39.7 Mb, 52% GC content), *Puccinia graminis* f. sp. *tritici* (176.7 Mb, 43.5% GC content) and *Saccharomyces cerevisiae* (12.2 Mb, 38.2% GC content). The number of small genome-encoded ORFs (≤ 600 bps) that are complete with a start and stop codon are 204,286 for *S. cerevisiae* genome, 537,720 for *Z. tritici* and 3,491,009 for *P. graminis* f. sp. *tritici*. The highest variability in GC content is observed in ORFs that encode very small proteins < 50 aas.

### EffectorGeneP annotates genes from transcripts with ML models and resolves false transcript fusions

Using known effectors as positive controls, we encountered recurring errors in a standard workflow of RNA-seq transcript assembly (Figure 2). Thus, we first optimized RNA-seq transcript assembly for effector discovery by assembling transcripts for each sample/replicate, merging these transcripts into a consensus set and re-predicting transcripts in genomic regions that show read coverage but did not have an assembled transcript. These transcript sets were then used as input for EffectorGeneP, which derives CDS candidates in each transcript and scores this with the ML models (Figure 3). To determine if the probability for a CDS is significant, EffectorGeneP uses 10,000 random CDS sequences with a similar codon content to the target genome to derive *z*-scores and *p*-values. If a CDS candidate has a significant probability, an overall gene score [0-100] using probabilities for their introns, UTRs and Kozak sequences is calculated and the highest-scoring gene model for the transcript is returned. Furthermore, EffectorGeneP returns the effector coding probability of a gene, which can be used downstream for ranking of candidates.

**Figure 2:**
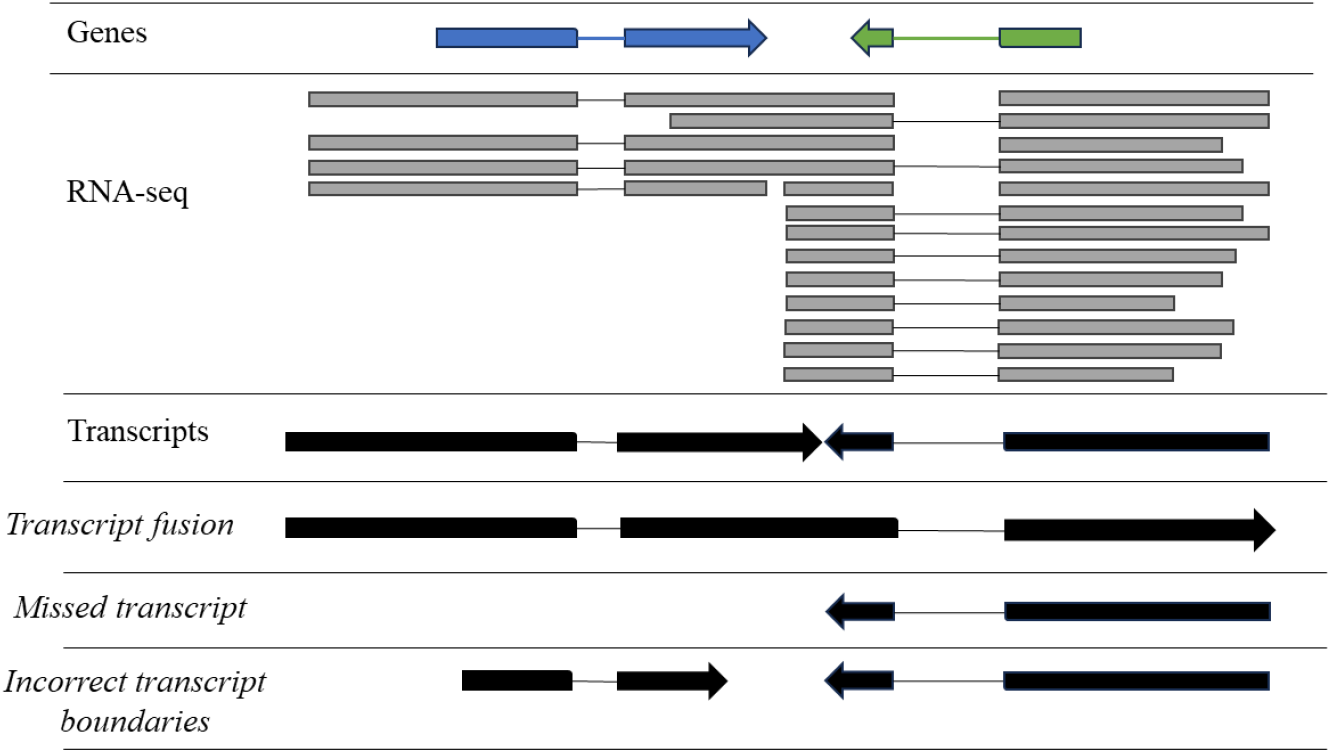
Commonly encountered errors during transcript assembly from RNA-seq reads. Two genes are shown, one in the forward and one in the reverse direction together with their RNA-seq read coverage. Each gene has one intron which can be seen in the RNA-seq read alignments. The correct transcript structures are shown, followed by commonly encountered transcript assembly errors. First, we frequently observed incorrect fusion of transcripts from adjacent genes into single larger transcripts, which would lead to missed or incorrect gene models under the one-transcript-encodes-one-gene hypothesis. Even with stranded RNA-seq data we observed that two opposing strands might be combined into one false fusion transcript with partially incorrect strand assignment. Second, we observed missed transcripts despite RNA-seq read support due to low coverage or overlapping UTRs with another transcript. Third, we encountered incorrect transcript boundaries due to low coverage which can lead to either truncated gene models (and potentially missed signal peptides) or missed gene models. Fourth, when using default parameters some transcript assembly methods will remove single-exon transcripts if they do not have high coverage.

**Figure 3:**
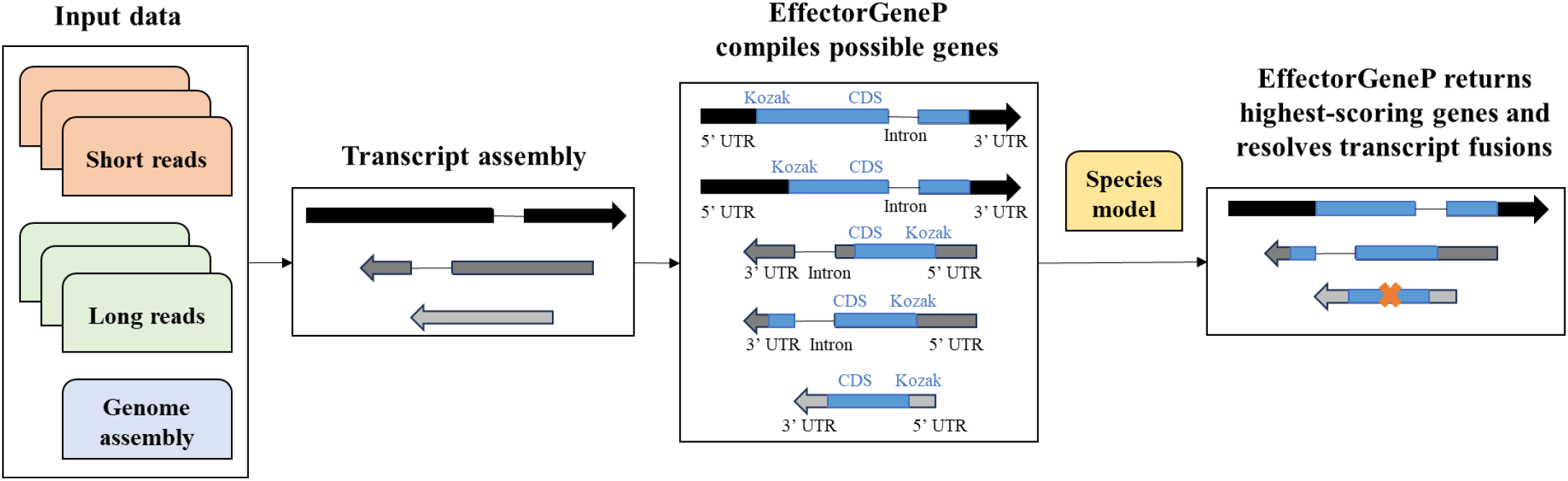
EffectorGeneP compiles gene candidates from transcripts and resolves transcript fusions. For a given transcript, EffectorGeneP compiles possible genes and scores their gene components (CDS, 3’ and 5’ UTRs, introns if present and Kozak sequences) with ML models. The highest-scoring gene is returned as the primary gene model and if long UTRs with low probability are present, additional gene candidates will be investigated in those UTRs to resolve transcript fusions. By default, EffectorGeneP will use the transcript strand information as its most likely direction of transcription, but it will also investigate gene candidates in the opposite direction. For each CDS, we also keep those smaller CDSs that have the same stop codon but a different start site. This allows selection of the highest-scoring translation start site as well as the detection of genes encoding secreted proteins where adjacent start sites might obscure the signal peptide region.

We illustrate how EffectorGeneP predicts the most likely gene per transcript for the *M. oryzae* effector gene *MC*69^20^, which encodes a small protein of only 55 aa. For the *MC*69 transcript, EffectorGeneP returns eight CDSs that encode proteins larger than 50 aa and could potentially constitute genes (Figure 4A, Supplementary Table 3). Candidate 6 (*MC*69) has the highest cumulative gene score with a high Kozak score and is returned as the best gene model, despite not being the longest CDS in the transcript. As a second example, we show the *U. maydis* effector gene *Eff1-2* which is in a transcript that has CDSs with the same stop codon but with three possible start sites in proximity (171 bp) (Figure 4B). The longest gene candidate encodes a protein of 506 aa, whereas the shortest gene candidate encodes the Eff1-2 protein with 449 aa. In the reference annotation, Eff1-2 was originally annotated as a longer, non-secreted protein but it was later manually corrected^21^. EffectorGeneP assigns a higher gene score to Eff1-2 than to the longest CDS and returns a high Kozak sequence score only for Eff1-2 (Supplementary Table 4). Again, EffectorGeneP selects Eff1-2 as the final gene model, despite it again not being the longest CDS in the transcript. As a third example, we correctly annotate the *M. oryzae* metallothionein 1 *(MMT1)* gene, which encodes a very small protein of 23 aa that is required for pathogenicity^22^. We ran EffectorGeneP with a minimum required protein length of 20 aa instead of 50 aa. For the *MMT1* transcript, EffectorGeneP returns eight CDSs which encode proteins of at least 20 aa (Figure 4C). However, only two of the eight have a significant coding probability and the *MMT1* gene has the highest score despite its very small length (Supplementary Table 5).

**Figure 4:**
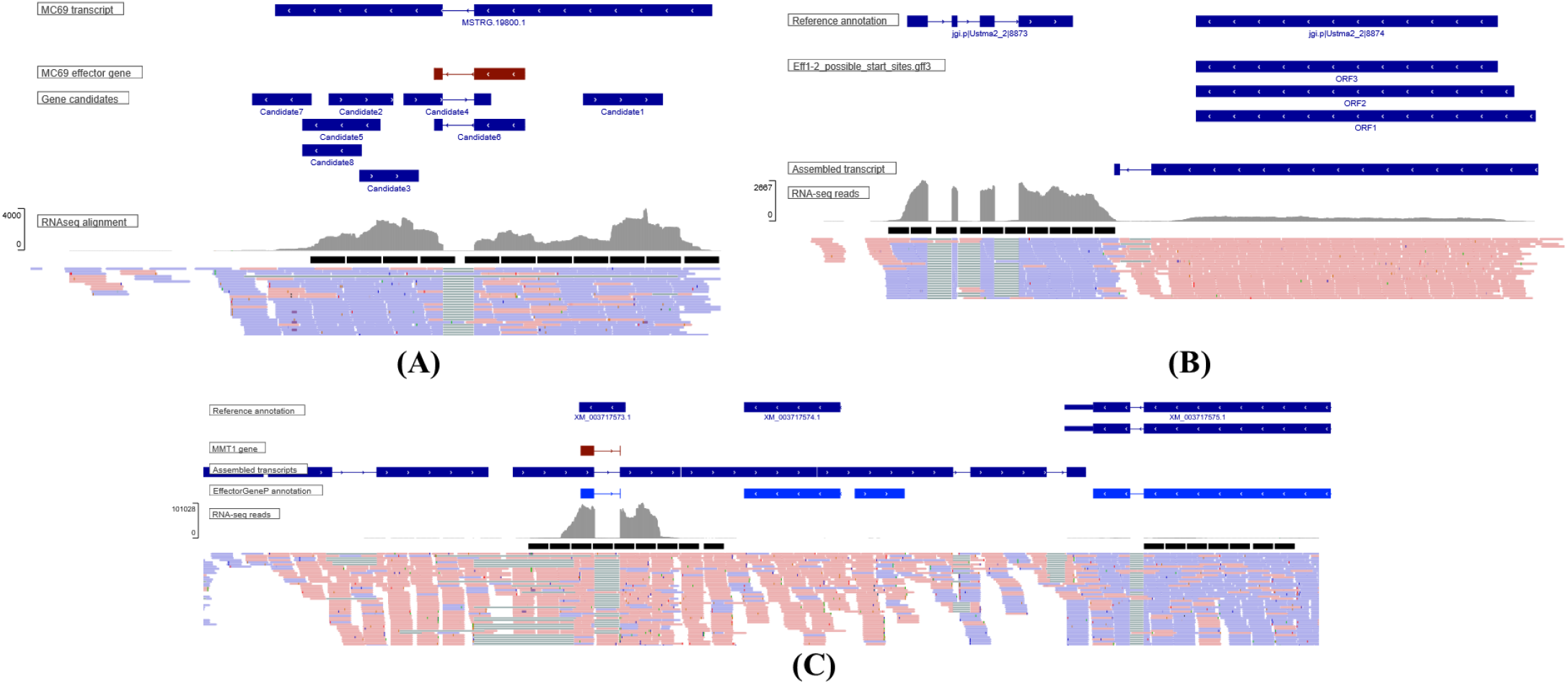
Selected examples for EffectorGeneP effector annotation.. **(A)** The assembled region that covers *MC*69 is shown with the majority of RNA-seq reads indicating transcription from the reverse strand. Four candidates CDS are present (four on the reverse strand, four on the forward strand). The *M. oryzae MC69* gene encodes a small protein of 55 amino acids and is not the longest ORF in the transcript, yet it is picked by EffectorGeneP as the highest-scoring gene. **(B)** The *U. maydis* effector gene *Eff1-2* is located in a transcript that has gene candidates with the same stop codon but with three possible start sites in close proximity. EffectorGeneP returns *Eff1-2* as the annotated gene for the transcript. **(C)** When setting the minimum protein length to 20 aas, EffectorGeneP correctly annotates the *M. oryzae* metallothionein 1 gene, which encodes a very small protein of 23 aas.

We observed numerous cases of false transcript fusions even with stranded RNA-seq data by manual inspection of transcript assemblies. EffectorGeneP addresses false transcript fusion events by investigating the 3’ and 5’ UTRs of the highest-scoring gene if the probability for being a UTR is low and they are sufficiently long to harbour additional genes (Supplementary Figure S1). This allows EffectorGeneP to correctly annotate multiple genes from false fusion transcripts whilst obeying the exon-intron structure of the transcripts. For example, the effector gene cluster in *U. maydis* that contains the seven consecutive genes *Eff1-5* to *Eff1-11* has three mis-assembled transcript fusions (Supplementary Figure S2). EffectorGeneP correctly resolves these false transcript fusions and annotates all effector genes in this cluster (Supplementary Figure S2).

**Supplementary Figure S1:**
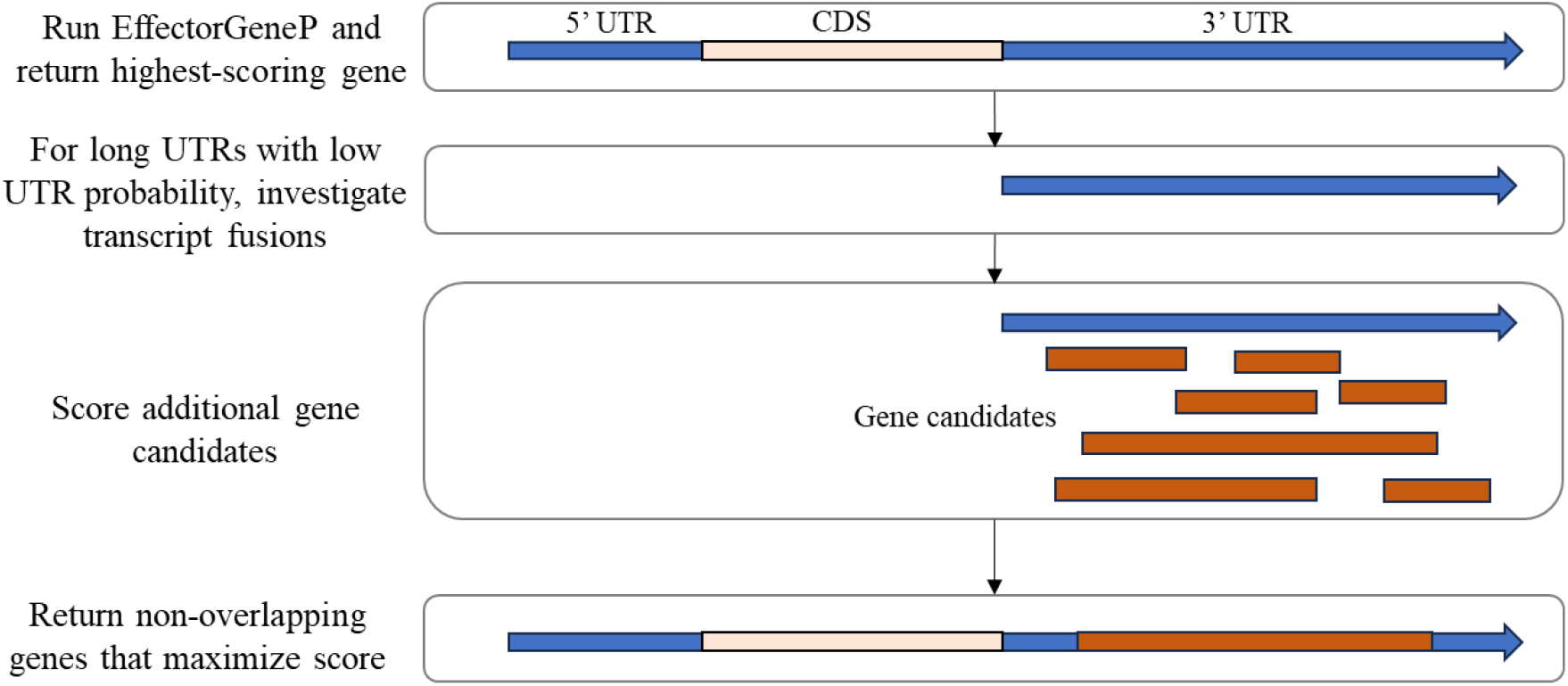
EffectorGeneP is able to resolve transcript fusions by investigating the 3’ and 5’ UTRs of the highest-scoring gene if the probability for being a UTR is low and they are sufficiently long to harbour additional genes. Each additional gene candidate is described as a genomic interval that has a weight (length-adjusted gene score). A weighted interval scheduling approach on the set of candidate genes returns the gene set that maximizes the overall score such that the final gene set is mutually compatible.

**Supplementary Figure S2:**
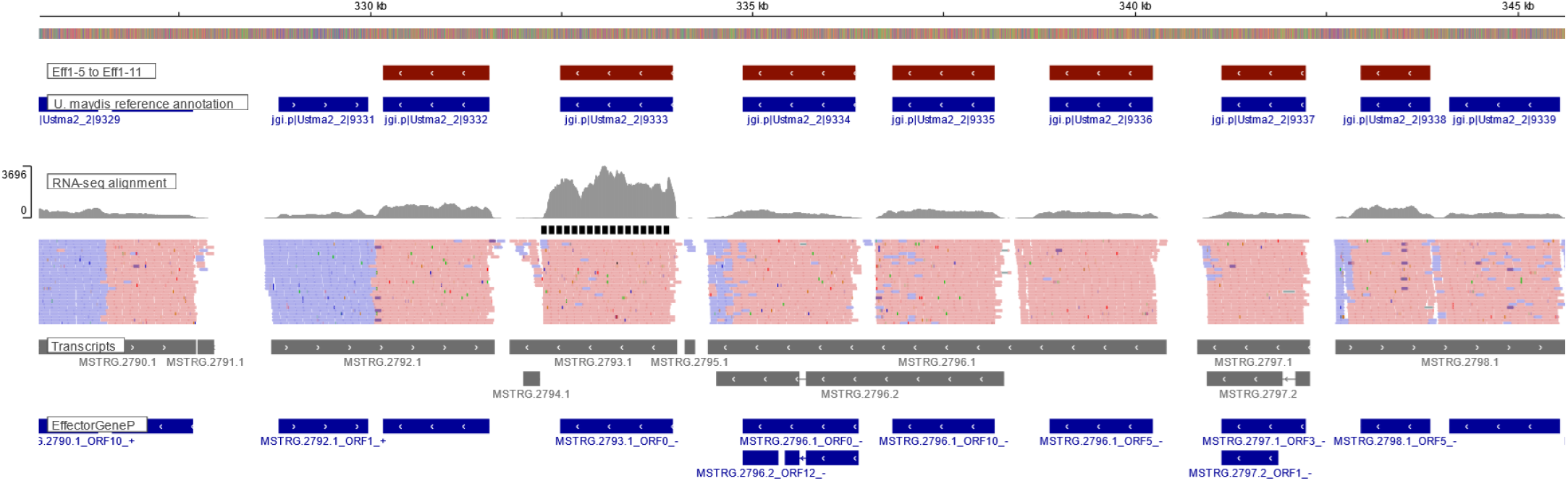
The *Ustilago maydis effector cluster Eff1-5* to *Eff1-11* has multiple transcript fusions. Eff1-5 (protein ID 9332) is an incorrect fusion of two transcripts on opposite strands. Second, the transcripts coding for Eff1-7 to Eff1-9 (protein IDs 9334-9336) are merged into a false fused transcript. Third, the transcript encoding Eff1-11 (protein ID 9338) is an incorrect fusion of two transcripts and additionally is assigned the incorrect strand by the transcript assembler.

### EffectorGeneP outperforms other gene prediction methods for effector gene annotation

We evaluated state-of-the-art gene annotation tools on a set of positive controls consisting of 173 known effector genes from ten genomes (Supplementary Table 6). First, we assessed if the 173 effectors are present in the set of transcripts assembled from the infection RNA-seq data sets (Supplementary Table 1), as these are the sole input for annotation by EffectorGeneP and TransDecoder. On average, the exon-intron structures of the assembled transcripts correctly assign 98% of the coding bases and 95% of the intron bases (Table 1), suggesting that most effector genes could be annotated directly from *in planta* RNA-seq data. Next, we compared the EffectorGeneP annotation to several other gene annotation methods that use either hybrid transcript/*ab initio* or purely *ab initio* approaches (Table 2). EffectorGeneP annotated 95% of the 173 known effector genes correctly, followed by CodingQuarry with 78% and Funannotate with 63.1% whereas BRAKER3, Helixer and TransDecoder only annotated 42%, 49% and 60% correctly, with high variation between the species (Table 2). BRAKER3 annotated none of the *P. graminis* f. sp. *tritici* and *P. nodorum* effectors and Helixer performed poorly for *P. graminis* f. sp. *tritici* and *Z. tritici*. One effector was missed by all methods expect EffectorGeneP (*AvrP4* in *M. lini*) and for two effectors a different protein was annotated at the locus by all methods including EffectorGeneP (*BAS162* and *MoCDIP5* in *M. oryzae*). In addition, we evaluated the output of several other methods included as subcomponents of Funannotate, however these all delivered poor results (AUGUSTUS: 35.7%; GeneMark: 37%; PASA: 49%; GlimmerHMM: 25%; SNAP: 15.1%). Taken together, this indicates that a standard gene annotation pipeline is likely to miss a substantial part of the effector complement of these pathogens.

**Table 1.**
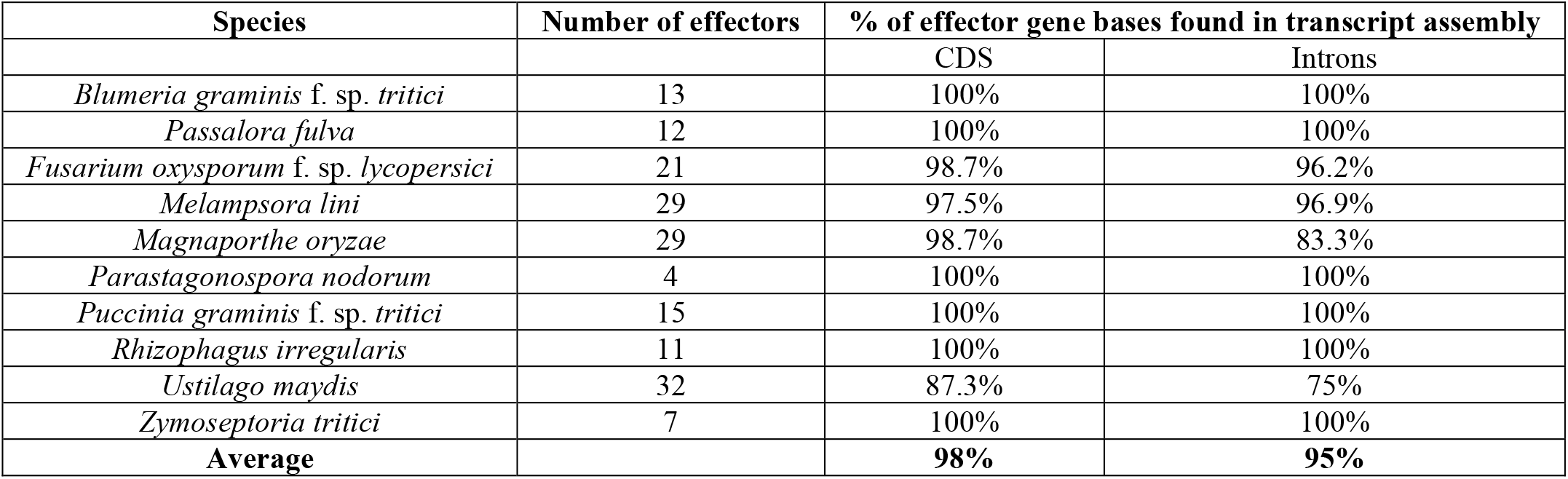
Base-level sensitivity for coding and intron sequences of known effector genes in the assembled transcripts.

**Table 2.** Benchmarking results for known effector genes of 10 fungal species. EffectorGeneP achieved the highest or equal highest proportion of correct effector gene annotations in every species. We used a stringent approach requiring a perfect match to the CDS of the known effector gene. In addition to EffectorGeneP, we ran CodingQuarry in pathogen mode (CodingQuarry-PM), providing the exact same set of input transcripts. We also ran the Funannotate pipeline, which wraps nine tools to achieve a consensus annotation, as well as Helixer, BRAKER3 and TransDecoder.

|  | Number of effectors | Correctly annotated effector genes |  |  |  |  |  |
| --- | --- | --- | --- | --- | --- | --- | --- |
|  |  | EffectorGeneP | CodingQuarry-PM | Funannotate | BRAKER3 | Helixer | TRANS-DECODER |
| *B. graminis* f. sp. *tritici* | 13 | **100%** | 84.6% | 92.3% | 69.2% | 69.2% | **100%** |
| *P. fulva* | 12 | **100%** | 75% | 50% | 58.3% | 33.3% | 33.3% |
| *F. oxysporum* f. sp. *lycopersici* | 21 | **90.5%** | 81% | 47.6% | 23.8% | 33.3% | 38.1% |
| *M. lini* | 29 | **89.7%** | 51.7% | 72.4% | 27.6% | 58.6% | 79.3% |
| *M. oryzae* | 29 | **89.7%** | 82.8% | 75.9% | 62.1% | 75.9% | 51.7% |
| *P. nodorum* | 4 | **100%** | **100%** | 50% | 0% | 25% | 75% |
| *P. graminis* f. sp. *tritici* | 15 | **100%** | 80% | 66.7% | 0% | 13.3% | 60% |
| *R. irregularis* | 11 | **100%** | 81.8% | **100%** | **100%** | **100%** | 81.8% |
| *U. maydis* | 32 | **84.4%** | 71.9% | 59.4% | 50% | 65.6% | 50% |
| *Z. tritici* | 7 | **100%** | 71.4% | 14.3% | 28.6% | 14.3% | 28.6% |
| **Average** |  | **95%** | **78%** | **62.9%** | **42%** | **48.9%** | **59.8%** |

### EffectorGeneP increases annotated secretome and effectorome sizes by 50%

After evaluation on known effector genes, we compared the EffectorGeneP annotations to the publicly available reference gene annotation. Where possible, we included sequencing data from both *in vitro* and *in planta* stages (Supplementary Table 1) as input for EffectorGeneP annotation to obtain close-to-complete secretome annotations. For several of the pathogens, EffectorGeneP led to a substantial increase in secretome size (on average 52% larger) and predicted effectorome sizes (on average 56% larger) (Table 3). The EffectorGeneP secretome of *P. fulva* was more than twice as large as the reference annotation secretome. In contrast, the secretome sizes of *F. oxysporum* f. sp. *lycopersici* and *S. cerevisiae* were in line with the reference annotations. For *P. nodorum* we only observed an increase of about 13% in the predicted secretome and effector proteins which can be explained by the high quality of the reference annotation, which was initially produced with CodingQuarry and underwent extensive manual annotation of gene models^23^. We also searched for sequence homologs of known effectors in the annotations. For 6 out of 10 species the EffectorGeneP annotations included more effector homologs than the reference annotations (e.g. 62 effector homologs in *U. maydis* compared to 38 in its reference annotation), suggesting that additional true effectors are being annotated. Because EffectorGeneP only annotates genes with transcript evidence, this is not due to annotation of non-expressed pseudogenes.

**Table 3:** Secretome sizes and predicted effectors for 11 fungal species. EffectorGeneP and reference annotations are shown. ** indicates a diploid genome. EffectorGeneP was run in conservative mode in this evaluation.

|  | EffectorGeneP annotation |  |  |  | Reference annotation |  |  |  |  |
| --- | --- | --- | --- | --- | --- | --- | --- | --- | --- |
| Species | # annotated genes \| gene isoforms encoding secreted proteins |  | # gene isoforms encoding secreted effectors | # genes with sequence homology to known effectors | # genes encoding secreted proteins | # genes encoding secreted effectors | # genes with sequence homology to known effectors | % increase of secretome \| effectorome |  |
| *B. graminis* f. sp. *tritici* | 2,088 | 2,275 | 1,317 (57.9%) | 283 | 1,268 | 802 (63.2%) | 289 | 79% | 64% |
| *P. fulva* | 2,655 | 2,832 | 1,140 (40.3%) | 14 | 1,200 | 433 (36.1%) | 10 | 136% | 163% |
| *F. oxysporum* f. sp. *lycopersici* | 1,841 | 1,907 | 754 (39.5%) | 26 | 2,183 | 784 (35.9%) | 15 | -13% | -4% |
| *M. lini* (**) | 5,298 | 6,062 | 3,046 (50.2%) | 43 | 3,319 | 1,654 (49.8%) | 39 | 83% | 84% |
| *M. oryzae* | 2,897 | 3,171 | 1,404 (44.3%) | 102 | 1,804 | 741 (41.1%) | 95 | 76% | 90% |
| *P. nodorum* | 1,985 | 2,106 | 727 (34.5%) | 8 | 1,865 | 642 (34.4%) | 10 | 13% | 13% |
| *P. graminis* f. sp. *tritici* (**) | 6,665 | 7,349 | 4,549 (61.9%) | 79 | 6,094 | 3,996 (65.6%) | 81 | 21% | 14% |
| *R. irregularis* (**) | 2,644 | 2,839 | 2,009 (70.8%) | 75 | 1,509 | 1,087 (72%) | 46 | 88% | 85% |
| *U. maydis* | 908 | 989 | 369 (37.3%) | 62 | 611 | 212 (34.7%) | 38 | 62% | 74% |
| *S. cerevisiae* | 272 | 272 | 112 (41.2%) | - | 279 | 108 (38.7%) | - | -3% | 4% |
| *Z. tritici* | 2,175 | 2,388 | 969 (40.6%) | 23 | 1,844 | 751 (40.7%) | 24 | 30% | 29% |

To further assess whether the additional annotated genes encode true protein products, we ran structure predictions using Boltz-2^24,25^ on both the EffectorGeneP and reference annotation secretomes of *B. graminis* f. sp. *tritici* and *M. oryzae*. The confidence values of the Boltz-2 structure predictions for the known effectors as well as sequence-related effector homologs ranged between 0.35-0.95 (mean 0.78, *n* = 573) for *B. graminis* f. sp. *tritici* and between 0.32-0.98 (mean 0.73, *n* = 198) for *M. oryzae* (Figure 5A). For *B. graminis* f. sp. *tritici*, we also extracted secreted proteins with an [YFW]xC or Y(x)xC sequence motif in their N-terminal region (*n* = 775). For these candidate effectors, the confidence values were slightly lower and ranged from 0.29-0.95 (mean 0.68). As confidence > 0.6 is generally considered to indicate a high-quality structure prediction, this implies that a significant proportion of effectors are modelled poorly (27% for *M. oryzae;* 13.4% for *B. graminis* f. sp. *tritici*; 36.6% for *B. graminis* f. sp. *tritici* [YFW]xC or Y(x)xC effectors). However, the low confidence values for many of the *bona fide* effectors suggests that this is not a good predictor of an incorrect gene annotation, especially if no sequence homologs or similar structures exist in databases. Thus, we investigated if the annotated secreted proteins fall into structurally-related families, possibly including known effector structures. We clustered the secretomes based on similarity in the predicted protein structures, including the available experimentally determined structures of *B. graminis* f. sp. *tritici, B. graminis* f. sp. *hordei* and *M. oryzae* effectors. We assigned structural clusters as ‘effector families’ if they contain a member with sequence homology to a known effector or if they form a cluster with a known effector structure. When considering structural effector family clusters with at least five members, EffectorGeneP leads to an 27.8% increase in members for *B. graminis* f. sp. *tritici* and an 30.6% increase for *M. oryzae* (Figure 5B). This strongly suggests that additional effector genes are captured in the EffectorGeneP annotations.

**Figure 5:**
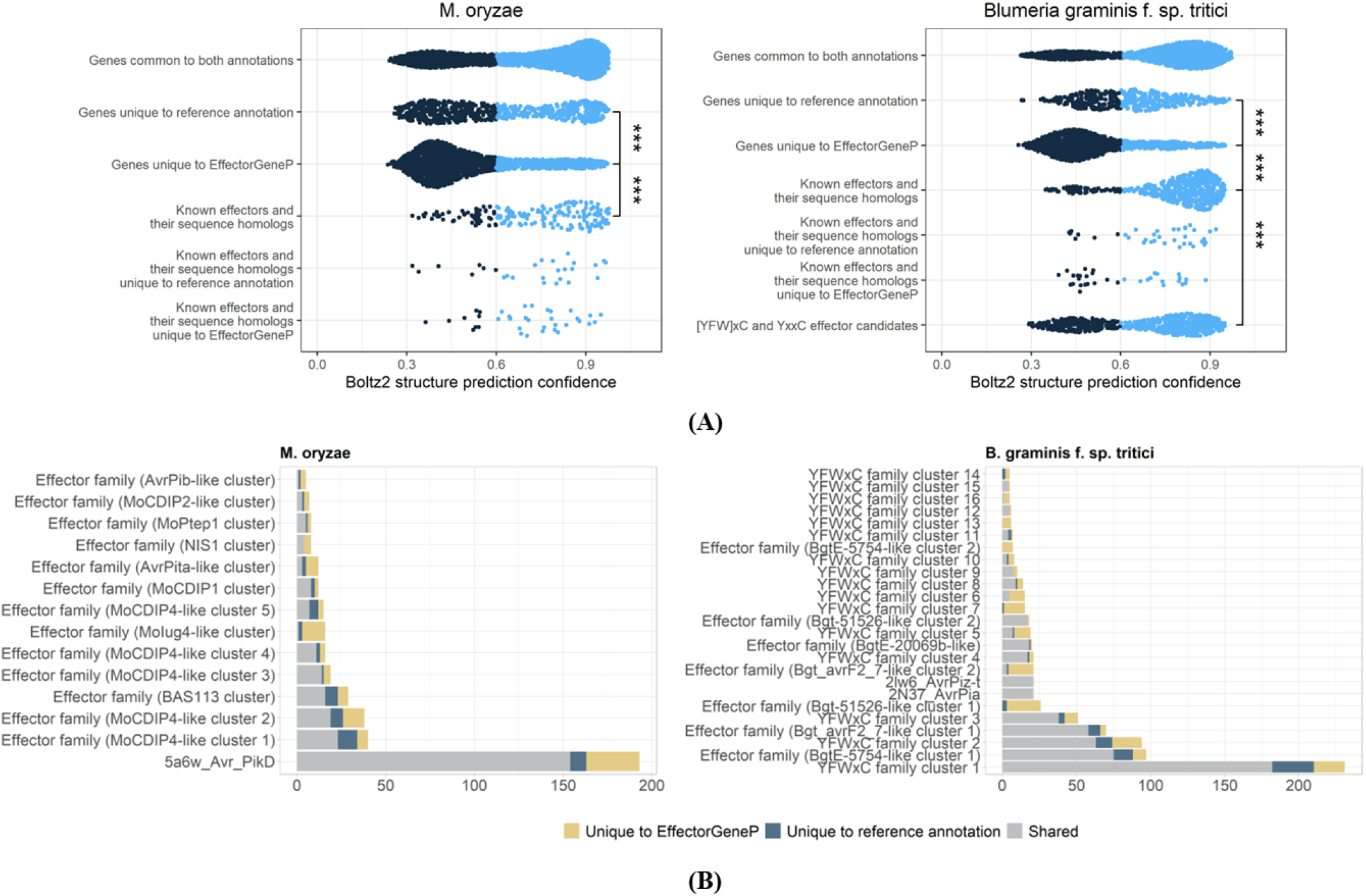
**(A)** Boltz-2 protein structure prediction confidence values for *M. oryzae* and *B. graminis* f. sp. *tritici* secretomes. Genes common to both annotations tend to have high confidence values, whereas the EffectorGeneP annotations and effector sets showed a number of proteins in the lower confidence range < 0.6. **(B)** Structural clustering of the predicted structures shows an expansion of genes annotated by EffectorGeneP in effector clusters.

### EffectorGeneP annotates a large family of *AvrSr26* effector genes encoding proteins only 30% sequence similarity

Accurate gene annotation is a pre-requisite for functional screening of effector candidates. We previously used the reference annotation of *P. graminis* f. sp. *tritici* isolate Pgt21-0^26^ to design and synthesise a library of 696 effector candidates, which was screened against several wheat stem rust resistance genes in protoplasts^27^. This approach was successful in identifying *AvrSr13* and *AvrSr22*, but no hits were identified with *Sr26*. This effector library was subsequently extended to 1,373 candidates^28^ after using an *ad hoc* process to expand the reference genome annotation^19^. Screening this expanded library with *Sr26* identified four effector candidates (clones 998, 1000, 1007 and 1033) that showed significantly reduced expression in wheat protoplasts in the presence of *Sr26* relative to an empty vector control (Figure 6A), induced cell death in the presence of *Sr26* in individual wheat protoplast assays (Figure 6B) and in *Nicotiana benthamiana* (Figure 6C, Supplementary Figure S3) or *N. tabacum* (Supplementary Figure S3). Furthermore, recombinant barley stripe mosaic virus (BSMV) expressing any of these four candidates was unable to infect the wheat line Avocet carrying *Sr26* (Figure 6D), confirming the function of these genes as *AvrSr26*. The four *AvrSr26* genes reside at separate loci and were designated as *AvrSr26*^*chr2*^*-01* (#998), *AvrSr26*^*chr11*^*-01* (#1000), *AvrSr26*^*chr12*^*-01* (#1007), and *AvrSr26*^*chr17*^*-01* (#1033) based on their chromosomal location. The allelic variant of *AvrSr26*^*chr2*^*-01* on chromosome 2B contains a frameshift mutation, while the *AvrSr26*^*chr12*^*-01* gene is identical in both alleles. The other two loci are heterozygous, with the allelic variants *AvrSr26*^*chr11*^*-02* and *AvrSr26*^*chr17*^*-02* encoding proteins containing amino acid sequence differences. Neither of these gene variants were in the screened effector library, but both induced cell death in wheat protoplasts when co-expressed with *Sr26* with *AvrSr26*^*chr17*^*-02* inducing an intermediate response (Figure 6B). Noticeably, the AvrSr26 proteins only share amino acid identity of ∼30% (Figure 6E). A sequence similarity search identified a large sequence-related family of 125 AvrSr26-like proteins in the EffectorGeneP annotation (124 genes with one additional isoform, amino acid identity: 23%), which also includes AvrSr13. In contrast, the CodingQuarry and Funannotate annotations only contain 105 and 60 AvrSr26-like proteins, respectively, some of which are not predicted to be secreted suggesting potential mis-annotation of their N-terminal regions (Supplementary Table 7). We combined the annotations, kept those that have a signal peptide and removed identical mature proteins which resulted in 141 unique AvrSr26-like protein sequences (Supplementary Table 8). A phylogenetic tree showed that the four recognized clones were not part of the same protein sub-family (Supplementary Figure S4). Lastly, we note that the Boltz-2 structure prediction confidence values of the 141 AvrSr26-like mature proteins were again quite variable, ranging from 0.32 to 0.85 (mean 0.61), with 36.2% of the proteins having low confidence of < 0.6. This demonstrates again that *bona fide* effector proteins can have poor structure predictions.

**Figure 6:**
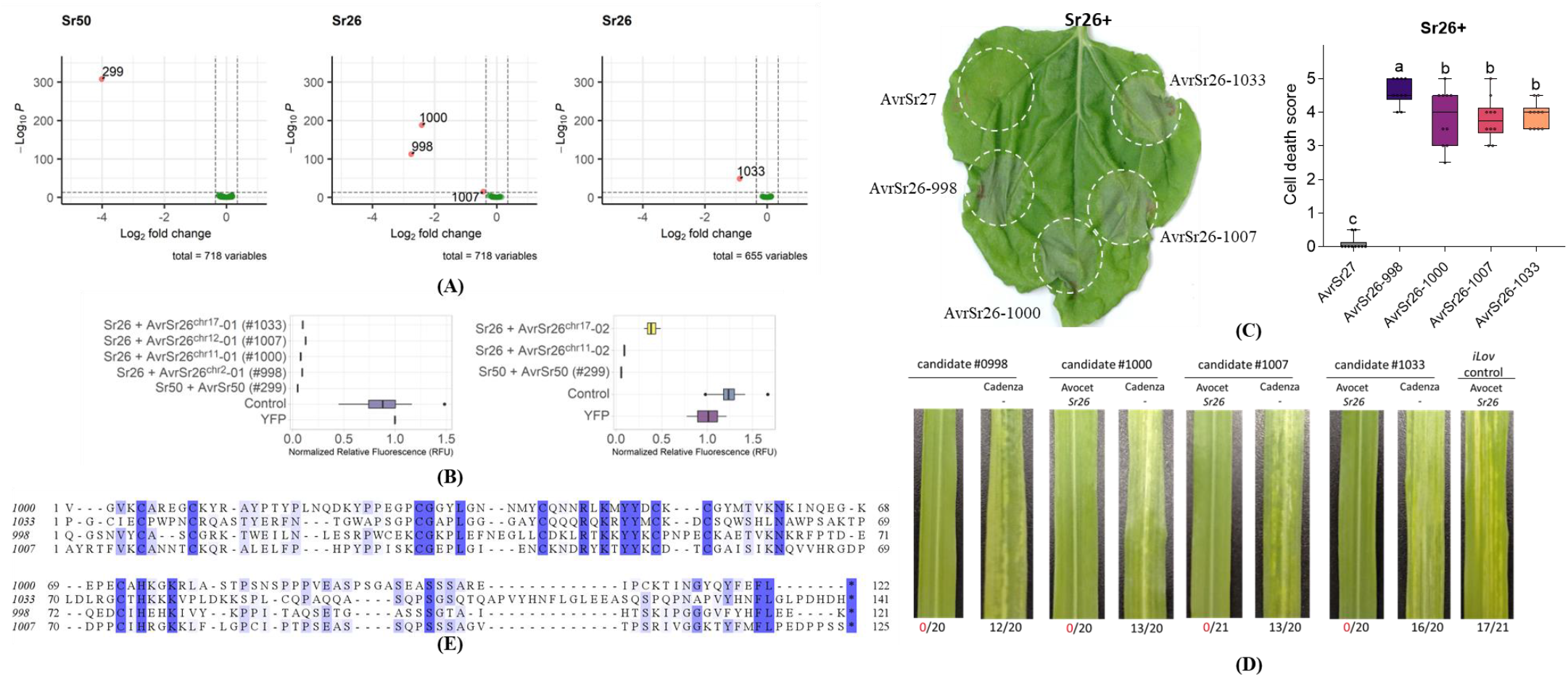
**(A)** *P. graminis* f. sp. *tritici* effector library screen with Sr26. Differential gene expression analysis of a pooled stem rust effector library co-transformed into wheat protoplasts with Sr50 and Sr26, each compared to the empty vector. Graph shows volcano plot of differential expression (X-axis) versus adjusted *p-*value (Y-axis) for each effector construct (dots). Effector gene constructs showing significantly reduced expression within each treatment are labelled with their library ID number, with AvrSr50 represented by clone 299. **(B)** *Sr26* and *AvrSr26* genes were co-expressed in wheat protoplasts and fluorescence of a co-transformed YFP reporter gene measured (RFU, x-axis). *Sr50* and *AvrSr50* were included as a positive control. Reduced YFP fluorescence when R and Avr are co-expressed is indicative of immune recognition resulting in cell death. Control refers to *Sr26* and the *AvrSr26* genes alone. **(C)** Co-expression assay in *Nicotiana benthamiana*. Dotted lines mark out the area infiltrated with an *Agrobacterium tumefaciens* suspension. Cell death score in all infiltrated *N. benthamiana* leaves represented as a box and whiskers plot derived from at least six replicate infiltration sites. The x-axis represents the different treatments, the y-axis represents the cell death score out of 5 (0=no cell death, 5=cell death over the whole inoculated area). Samples marked by identical letters in the plot do not differ significantly [P < 0.05; analysis of variance (ANOVA), Tukey post hoc test]. **(D)** BSMV VOX analysis of *AvrSr26* candidates in wheat. BSMV strains expressing the four *AvrSr26* candidates and a negative control strain expressing a small fluorescent flavoprotein iLov were used to infect wheat varieties Avocet (carrying *Sr26*; Yu et al. 2010) and Cadenza (not known to contain any stem rust resistance genes). Images show typical symptoms at 24 days post-inoculation, and the numbers below indicate the successful infections observed out of the total inoculated plants from two replicate experiments. **(E)** Alignment of the *AvrSr26* effector proteins with signal peptides removed.

**Supplementary Figure S3:**
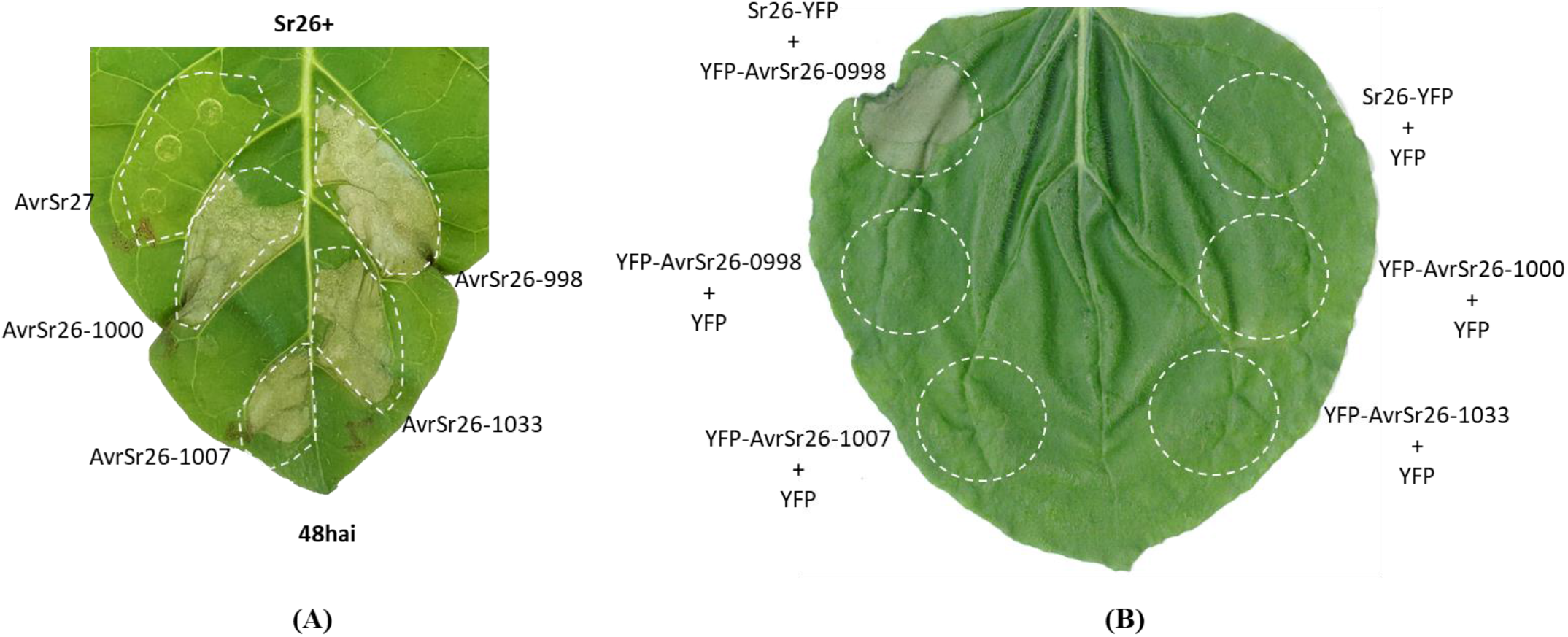
**(A)** Validation of the interaction between *Sr26* and the *AvrSr26* clones by co-expression in *Nicotiana tabacum* at 48 hai. Dotted lines mark out the area infiltrated with an *Agrobacterium tumefaciens* suspension. **(B)** Infiltrations of the *AvrSr26* clones without *Sr26* in *Nicotiana benthamiana* did not result in cell death.

**Supplementary Figure S4:**
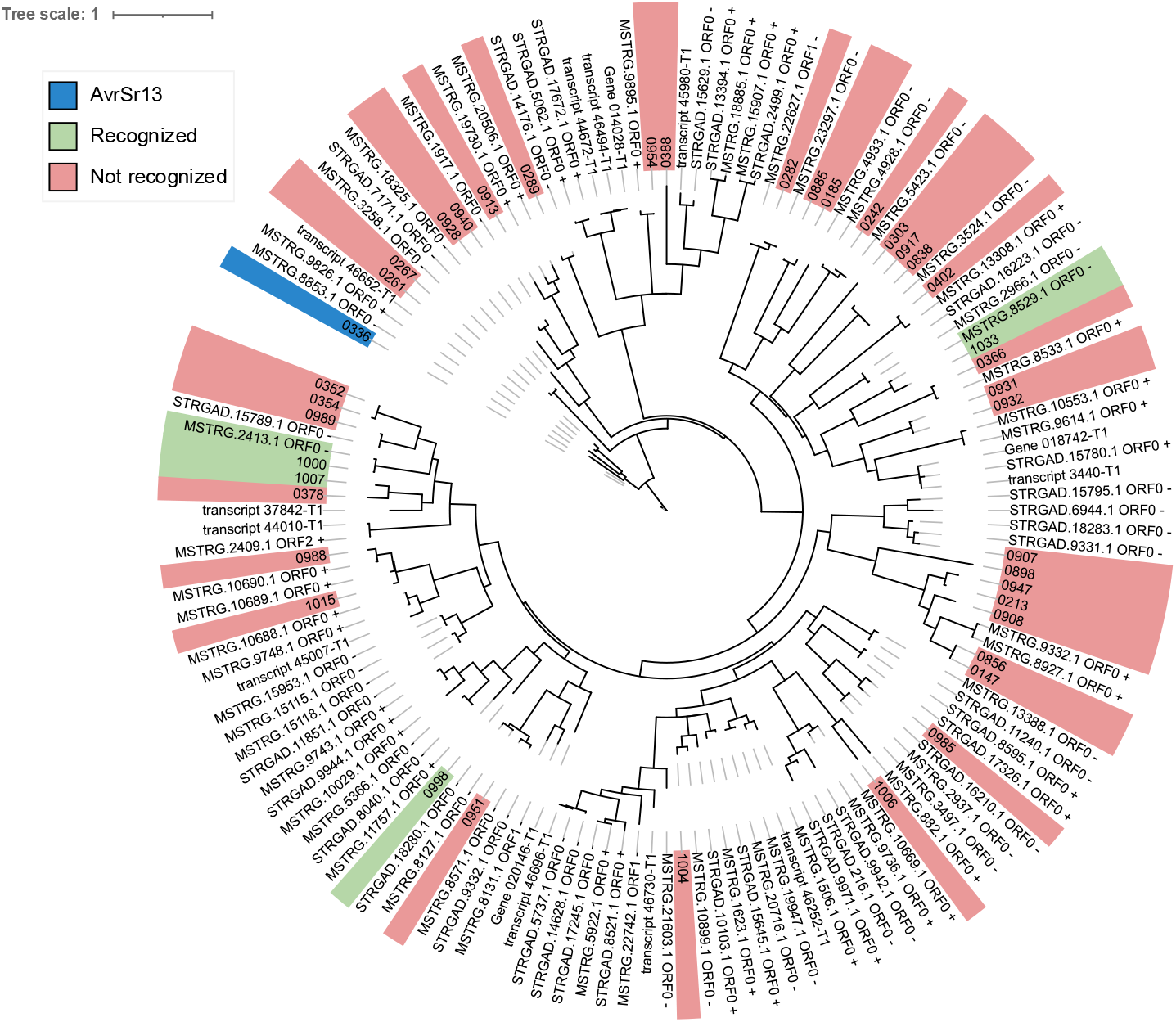
Phylogenetic tree of mature protein sequences for the AvrSr26-like family. Genes highlighted in red are present in the effector candidate libraries but were not detected as recognized by *Sr26* in wheat protoplasts.

### Species-specific models achieve higher prediction accuracy and higher effector coding scores

To understand why effector genes are difficult to annotate, we investigated which features are most discriminative in the ML classification for three pathogens: *P. graminis* f. sp. *tritici* with a highly repetitive genome (176.7 Mb, 43.5% GC), *M. oryzae* with a small genome and relatively high GC content (41 Mb, 51.6% GC), *Z. tritici* with similar genome size and GC content (39.7 Mb, 52.1% GC); as well as *S. cerevisiae* as a non-pathogen with a very small genome and very low GC content (12.2 Mb, 38.2% GC). For all three pathogens, we observed a significant difference in Codon Adaptation Index (CAI) for the CDSs of predicted effectors (Supplementary Figure S5). In *P. graminis* f. sp. *tritici*, we also noticed significantly lower GC content in CDSs of predicted effectors and in line with this an enrichment for A at the third position in codons. In contrast, *Z. tritici* has significantly higher GC content in CDSs of predicted effectors (Supplementary Figure S5) and a strong enrichment for C at the third position in codons. In *M. oryzae*, we found significantly lower GC content in CDSs of predicted effectors compared to secreted non-effectors as well as an enrichment for A and a depletion for G and T at the third position in their codons. In contrast, no significant difference in CAI was observed for the different classes of CDSs in *S. cerevisiae*.

Next, we tested the effect on effector gene annotation when using a different species model. Under the *P. graminis* f. sp. *tritici* model, all its effector genes were annotated correctly and attained very high EffectorGeneP scores of at least 0.89 (mean 0.99). When applying other species models to *P. graminis* f. sp. *tritici*, some of the effector genes failed to be annotated perfectly due to different start sites and overall, they had lower EffectorGeneP scores (Supplementary Figure S6). When using the model from the most closely related other pathogen, *M. lini*, the *P. graminis* f. sp. *tritici* effectors attained relatively high scores and were all annotated correctly. In line with the observation of GC content differences in effector CDSs (Supplementary Figure S5), the *Z. tritici* model performed poorly on *P. graminis* f. sp. *tritici* and rendered the EffectorGeneP scores ineffective for effector gene ranking. For *M. oryzae*, applying other species models also resulted in lower EffectorGeneP scores, with the lowest score of 0.17 recorded for *AvrPib* under the *Z. tritici* model and the *S. cerevisiae* model performing poorly with eight missed genes. Lastly, we investigated *U. maydis*, a species where only ∼28% of reference gene models have introns. *S. cerevisiae* is another species with mostly single-exon genes (∼95%), however the *S. cerevisiae* model performed poorly for *U. maydis* and did not call genes for five effector loci (Supplementary Figure S6). Taken together, this suggests that a species-specific or closely related species model will perform best.

**Supplementary Figure S5:**
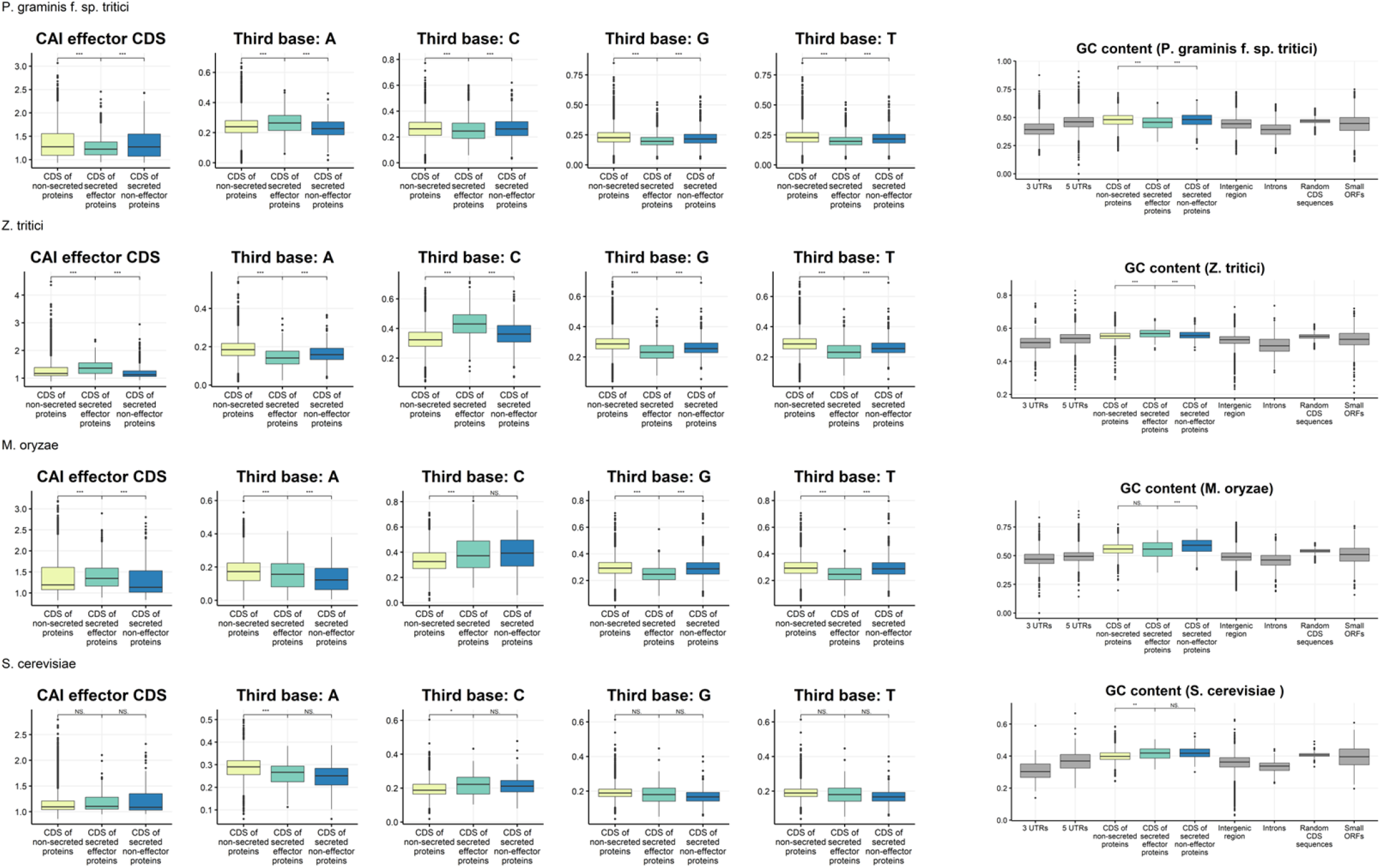
Codon adaptation index and distribution of third base in codons for three classes: CDS of non-secreted proteins, CDS of secreted effector proteins and CDS of secreted non-effector proteins. GC content for all classes of genomic features.

**Supplementary Figure S6:**
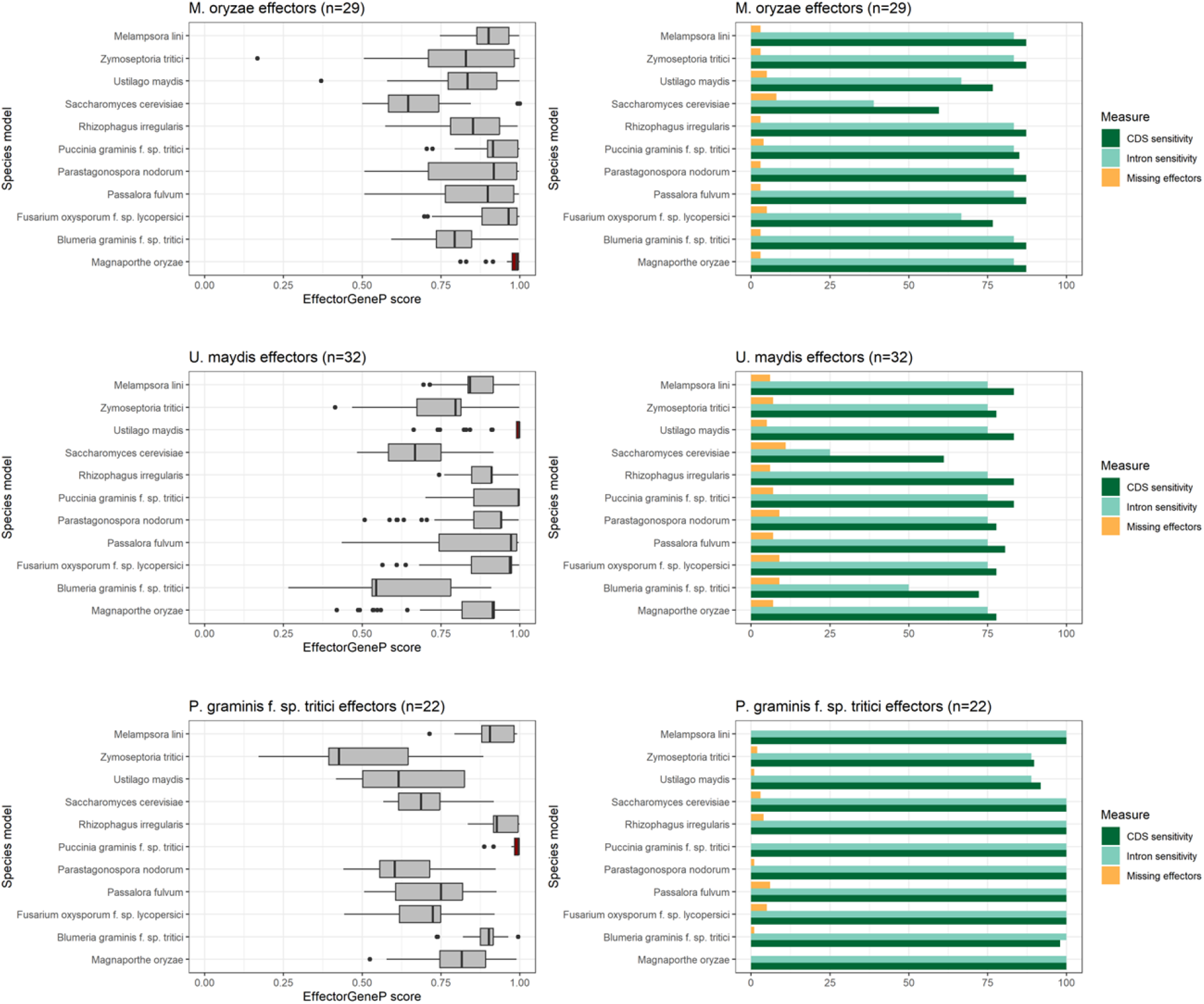
EffectorGeneP scores for known effectors under the different species models. Missing effectors are those that are not annotated perfectly.

### Using Iso-Seq data improves transcript continuity and isoform discovery, but might miss small effector genes

To test the effect of long-read transcript sequencing on annotation quality, we obtained both RNA-seq data and PacBio Iso-Seq data from infected leaves (6 dpi) for *P. graminis* f. sp. *tritici* strain Pgt21-0. Thus, differences in transcript assembly and gene annotation will reflect technical variation and not biological variation. Furthermore, we obtained PacBio HiFi genome sequencing data for the Pgt21-0 strain and re-assembled the genome, which resolved an additional 0.53 Mb of sequence for the 36 chromosomes and substantially reduced the number of gaps (Supplementary Table 9). We assembled transcripts for the RNA-seq data, the Iso-Seq data as well as from the combined RNA-seq/Iso-seq data using the hybrid transcript assembly mode of StringTie^29^ (Supplementary Table 10, Figure 7A). The EffectorGeneP annotations for these three sets contained 5,878 to 6,431 gene loci encoding secreted proteins with the highest number annotated using the hybrid transcripts and as expected, the highest number of isoforms annotated in the Iso-Seq transcripts (Supplementary Table 11). Notably, the RNA-seq and hybrid transcripts had the highest number of single-exon genes encoding secreted proteins. In line with this, the highest number of members of the single-exon *AvrSr26*-like family were annotated with the RNA-seq data (67.2%), whereas the Iso-Seq data only annotated 23.2%. The absence of two other effectors from the Iso-Seq annotation can be explained with a lack of raw read coverage of these genes. The Iso-Seq data did not show raw read coverage for the *avrSr27-3* gene and the *AvrSr62-6* gene, which both have low expression^28,30^. However, the *AvrSr26-*like genes that are missing from the Iso-Seq set do not have lower expression (Figure 7B). Alternatively, it is possible that small genes such as *AvrSr26* are under-represented in the Iso-Seq annotation which returned longer CDSs than the other two annotations (Figure 7C). Unexpectedly, despite high RNA-seq read coverage, StringTie failed to assemble AvrSr50 in both the RNA-Seq and hybrid mode (Supplementary Figure S7). Surprisingly, re-running the transcript assembly with the same parameters on only the RNA-seq reads that map to the *AvrSr50* gene region resulted in an assembled *AvrSr50* transcript. Taken together, whilst an improvement in overall gene annotation is to be expected with Iso-Seq data, it might currently miss some small effector genes when not complemented with RNA-seq data.

**Figure 7:**
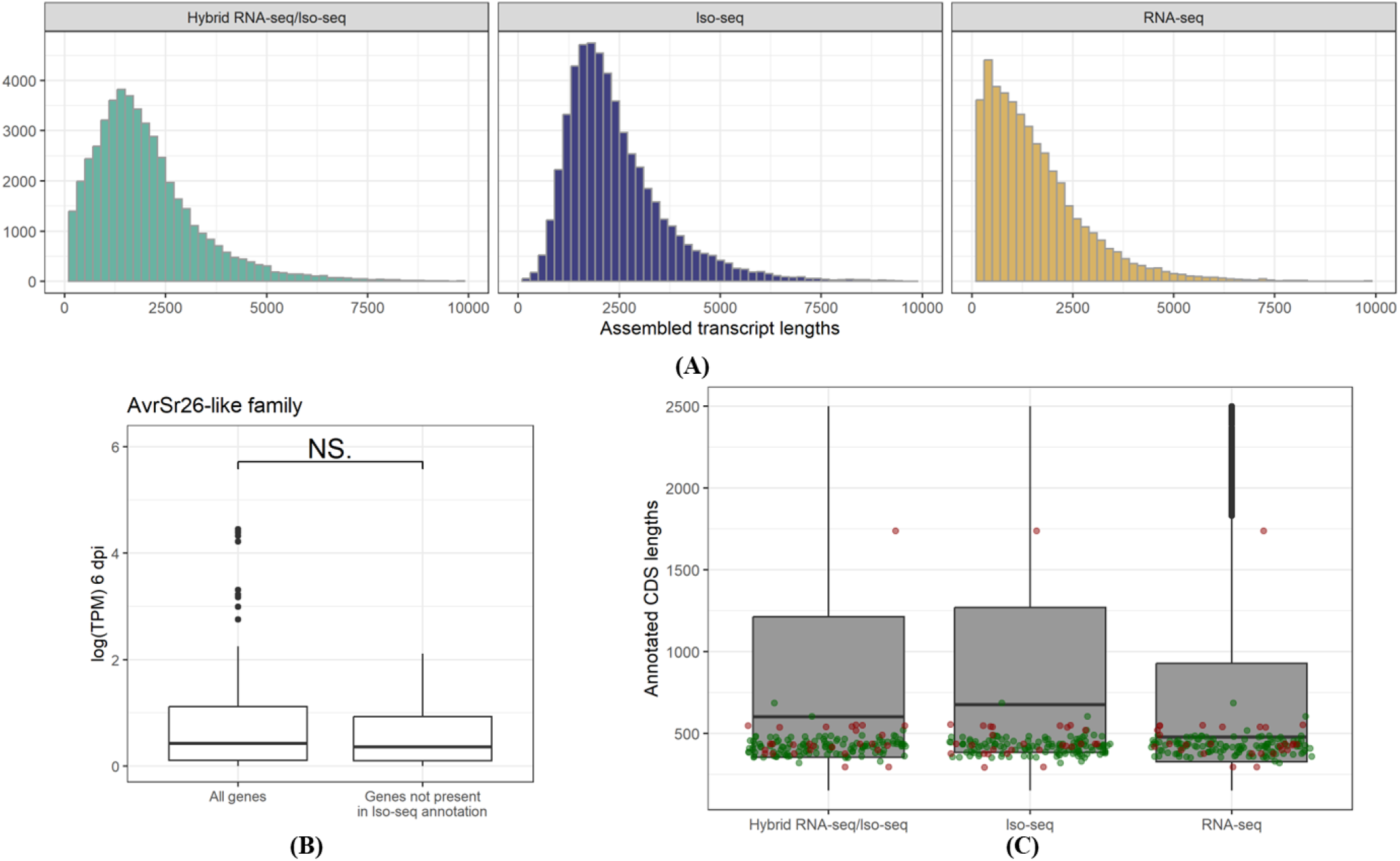
**(A)** Assembled transcript lengths for hybrid RNA-/Iso-Seq assembly, Iso-Seq assembly and RNA-seq assembly. The assembled transcript lengths are longer in the Iso-Seq and hybrid RNA-/Iso-Seq sets, which can be explained by potential fragmentation of transcripts in the RNA-seq assembly due to low coverage, bias of the Iso-Seq library preparation towards longer transcripts or enhanced resolution of UTRs in the Iso-Seq transcripts. **(B)** Expression at 6 dpi for *AvrSr26-like* genes not present in the Iso-seq annotation is not significantly different to all *AvrSr26-like* genes. **(C)** CDS lengths for three EffectorGeneP secretome annotations produced from hybrid RNA-/Iso-Seq transcript assembly, Iso-Seq transcript assembly and RNA-seq transcript assembly (red dots: known effectors and their alleles; green dots: AvrSr26-like family).

**Supplementary Figure S7:**
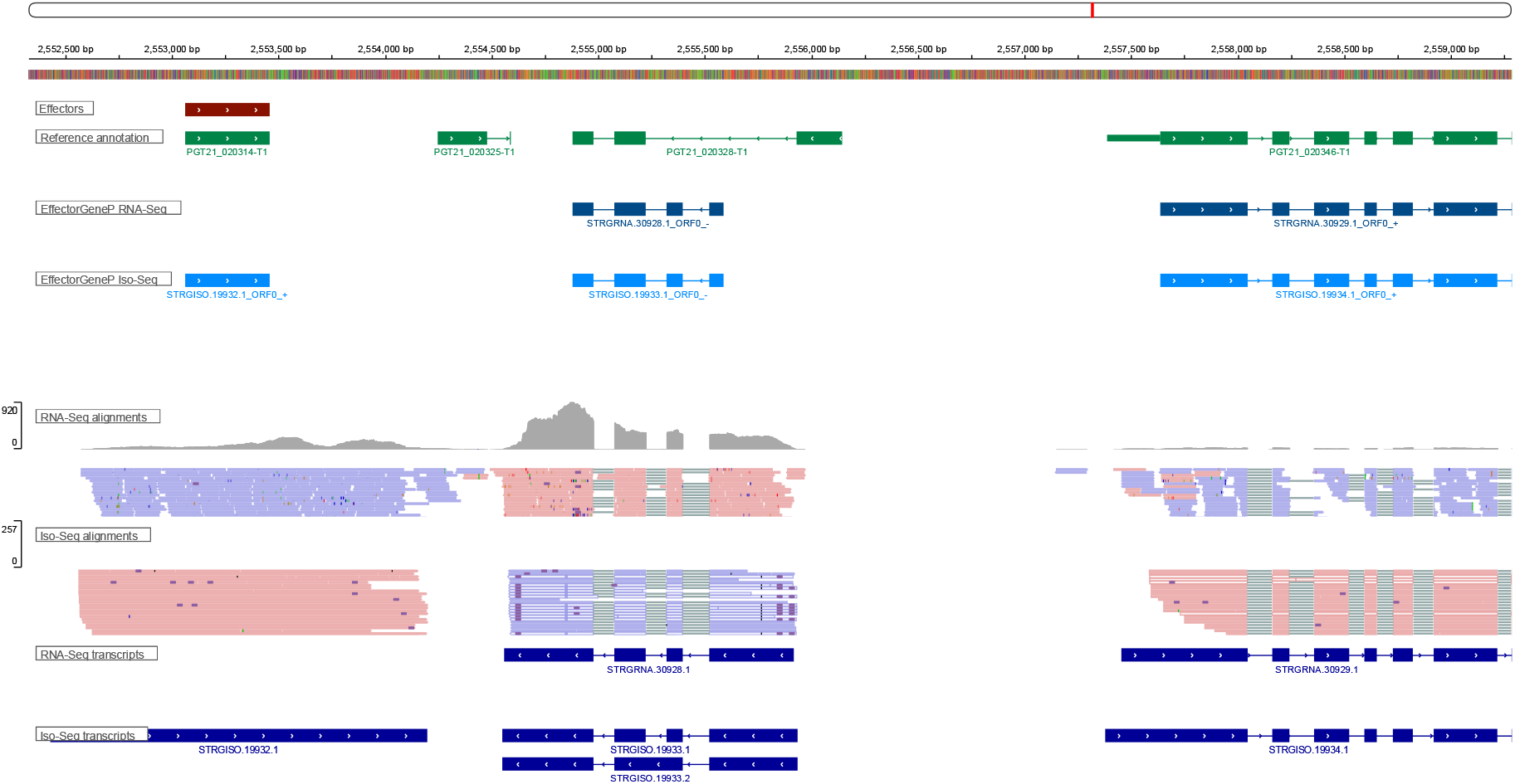
The AvrSr50 transcript (gene: PGT21_020314-T1) is missing from the RNA-seq assembly despite read coverage whereas the Iso-Seq data has an assembled transcript.

### Practical recommendations for using EffectorGeneP as part of a gene annotation pipeline

We have shown that EffectorGeneP is a useful tool for annotation of non-canonical genes such as effector genes. Routine practice in whole genome annotation is to integrate various annotation tools and select a consensus gene model at a particular locus. For instance, Funannotate uses EVidenceModeler^31^ with weights assigned to each of the its input sources. We tested providing the EffectorGeneP secretome annotation to Funannotate as an additional input source and found that this improved the annotation of known effector genes to a certain degree in the test genomes (Table 4). However, a stand-alone EffectorGeneP annotation achieves the highest sensitivity for these effector genes except in the case of *M. lini* where the non-expressed *AvrL2* gene variants are annotated by Funannotate but not with EffectorGeneP. Even when providing a very high weight to EffectorGeneP, some effector genes are lost in the EVidenceModeler process. Thus, for an application requiring sensitive effector gene annotation, such as for effector library design or a targeted effector search in a genomic interval, it would be preferable to use EffectorGeneP as a stand-alone tool and merge the EffectorGeneP secretome annotation with a reference annotation.

**Table 4:** Benchmark of a hybrid EffectorGeneP and Funannotate annotation. Default settings for Funannotate weights are Augustus: 1; Augustus High-Quality: 2, GeneMark: 1; GlimmerHMM: 1; Pasa: 6, SNAP: 1.

|  | Number of effectors | EffectorGeneP | Funannotate | Funannotate with EffectorGeneP (weight for EffectorGeneP) |  |  |  |
| --- | --- | --- | --- | --- | --- | --- | --- |
|  |  |  |  | 1 | 6 | 10 | 20 |
| *B. graminis* f. sp. *tritici* | 13 | **100%** | 92.3% | **100%** | **100%** | **100%** | **100%** |
| *P. fulva* | 12 | **100%** | 50% | 50% | 83.3% | 91.7% | 91.7% |
| *F. oxysporum* f. sp. *lycopersici* | 21 | **90.5%** | 47.6% | **90.5%** | **90.5%** | **90.5%** | **90.5%** |
| *M. lini* | 29 | **89.7%** | 72.4% | 72.4% | 72.4% | 93.1% | 93.1% |
| *M. oryzae* | 29 | **89.7%** | 75.9% | 79.3% | **89.7%** | **89.7%** | **89.7%** |
| *P. nodorum* | 4 | **100%** | 50% | 75% | 75% | 75% | 75% |
| *P. graminis* f. sp. *tritici* | 15 | **100%** | 66.7% | 66.7% | **100%** | **100%** | **100%** |
| *R. irregularis* | 11 | **100%** | **100%** | **100%** | **100%** | **100%** | **100%** |
| *U. maydis* | 32 | **84.4%** | 59.4% | 68.8% | 81.3% | **84.4%** | **84.4%** |
| *Z. tritici* | 7 | **100%** | 14.3% | 42.9% | 85.7% | **100%** | **100%** |
| **Average** |  | **95%** | 63% | 75% | 88% | 92% | 92% |

## Discussion

EffectorGeneP distinguishes itself from other annotation tools as follows. First, it is a transcription-based annotation tool that uses ML models instead of GHMMs, the ML models are trained separately on different classes of genes and the gene scoring process mimics the manual annotation process from transcription data that a human might follow. Second, EffectorGeneP learns gene properties from transcriptional data alone which allows for annotation of non-canonical genes such as orphan genes, small genes, single-exon genes or genes with non-canonical splice sites or spliced start/stop codons. Third, a dedicated effector coding model enables the accurate annotation of this class of genes. This model also returns an effector-coding probability for each gene, which could be used to prioritize effector candidates for functional analyses. Different species models are interchangeable to a certain degree, which might be determined by the GC content of the species but also other factors such as codon usage bias^32^. If using the EffectorGeneP scores for ranking, this should only be done using the ML models of the species.

We identified the accuracy of the assembled transcripts as the main limitation of EffectorGeneP. It would be a simple process to combine the transcript assembly with protein homology evidence and provide this combined set to EffectorGeneP. A future extension could also be the prediction of transcribed regions from RNA-seq alignments through ML models. The presence of pre-mRNA or genomic DNA contamination in RNA-seq data can be problematic for transcription-based annotation tools. We also noticed that possibly due to the dense gene space in fungal genomes and the enrichment of effectors for single-exon genes, refinements of standard transcript calling workflows might be necessary. For example, despite high coverage a two-round procedure of transcript calling was necessary to capture the AvrSr50 effector gene, which is located in a single-exon transcript adjacent to a multi-exon transcript. Whilst protein structure prediction tools have been suggested to be promising for indicating the quality of gene annotations^33^, we showed that a substantial proportion of *bona fide* effector proteins have low confidence structures, which can be explained by the lack of homologs in databases or by the presence of intrinsically disordered regions as described in some bacterial and oomycete effectors^34,35^. Currently, a low confidence structure prediction should not lead to the exclusion of a gene model when annotating effectors.

We demonstrated the use of EffectorGeneP for the discovery of the *P. graminis* f. sp. *tritici* effector *AvrSr26*. The *Sr26* resistance gene was introgressed into wheat from *Thinopyrum ponticum* and provides resistance to all tested wheat stem rust isolates including recent epidemic strains from Africa and Europe^36^. Given the deployment of *Sr26* in Australian wheat varieties for over 50 years, this represents a remarkable level of durability^37,38^. This durability can now be explained by the finding that *Sr26* recognises effectors from at least four genetically independent *Avr* loci, meaning that virulence evolution would require multiple independent mutations. This is effectively equivalent to a pyramid of four different resistance genes that each recognise a single *Avr* locus. Thus, it is likely that *Sr26* will continue to provide effective and durable resistance, especially when deployed in the context of the smaller recombinant segment that also contains another broadly effective resistance gene, *Sr61* ^39^. Such direct recognition of multiple effectors by a single NLR is very rare. Kunz *et al*. ^40^ recently found that the *Pm3e* resistance gene in wheat recognises two unrelated effectors from wheat powdery mildew, while WRR4A and WRR4B in *Arabidopsis thaliana* recognise eight and four diverse CCG class effectors from *Albugo candida*, respectively ^41^. Structural analysis indicates that WRR4A interacts with common peptide-backbone features of these sequence-diverse but structurally related effectors^42^. Given that only 41 out of 141 *AvrSr26*-like genes were included in the effector library screen, it is possible that additional members of this expanded gene family are also recognised by *Sr26*.

Rather than focussing on annotation of effector genes, EffectorGeneP could be re-trained on other classes of genes whose features may deviate from those of conserved genes. For instance, genes encoding small secreted, cysteine-rich peptides in plants have been shown to be missing from annotations^43^. It is not always obvious as to why a class of genes is difficult to annotate, however the most likely cause is their under-representation in the set used for GHMM training and the use of only one GHMM for the diverse set of genes present in a genome. The increased use of ML for gene annotation together with other high-quality sources of data will continue to advance accurate gene discovery across diverse species.

## Online Methods

### Curation of EffectorGeneP training sets

EffectorGeneP needs the following inputs for training of the ML models: 1) a genome assembly and 2) transcriptional data such as RNA-seq or Iso-Seq from infection stages. For training, we aligned the RNA sequencing reads against the genome assembly as follows: reads were cleaned with fastp 0.23.2 and default settings^44^, replicates/samples were concatenated and aligned with STAR 2.7.9a in 2-pass mode (-alignIntronMin 5 --alignIntronMax 3000 -- alignMatesGapMax 3000 --outFilterMultimapNmax 100) which removes junctions supported by <= two reads^45^. Transcripts were assembled with Stringtie 2.2.1 (-s 1 -m 200) with the appropriate strand setting^46^. EffectorGeneP then derives the following training sets from the resulting set of transcripts. First, for each transcript the longest ORF is extracted and translated into a protein. All longest ORFs are homology-reduced (cd-hit -c 0.8)^47^ and separated into three sets: 1) CDSs encoding non-secreted proteins (including transmembrane proteins with a signal peptide); 2) CDSs encoding predicted secreted non-effector proteins; and 3) CDSs encoding predicted secreted effector proteins. Secreted proteins are predicted as those that have a signal peptide (SignalP 4.1 -u 0.34 -U 0.34) and no transmembrane domains outside the N-terminal signal peptide region (TMHMM 2.0)^48,49^. Effectors are predicted with EffectorP 3.0^50^. From these CDSs and transcripts, we derive introns and UTRs and kept those with a minimum length of 6 bps (maximum length for UTRs: 3000 bps) as well as Kozak sequences (defined as the sequence starting nine bases before the start codon and ending one base after the start codon). We also derived intergenic regions (minimum length of 50 bps and maximum length of 3000 bps) as those sequences in the genome that are not covered by the transcripts. We removed homology in the intron, UTR and intergenic sequence sets with bbmap (dedupe.sh absorbrc=t minidentity=80 minoverlappercent=80) (http://sourceforge.net/projects/bbmap/). Introns or intergenic sequences that contain N’s were also removed. We predicted small, complete ORFs (<= 600 bps) with a start (ATG) and stop codon (TAA, TAG, TGA) from the full genome sequence^51^. We removed ORFs that are similar to the CDS of infection-expressed transcripts (cd-hit-2d -c 0.9 -n 5 -d 0)^47^ to remove redundancy between training sets. We also removed small ORFs that are predicted to be secreted (same method as described above) as these could constitute *bona fide* single-exon effector genes. We used compseq^52^ to calculate expected trinucleotide and hexanucleotide frequencies for the training sets. We also produced 10,000 random CDS sequences by randomly sampling codons from the small ORFs and sequence lengths from the CDSs of the longest ORFs. Lastly, we generated 10,000 random Kozak sequences for training.

### Training of a ML classifier for genomic features

For training of the ML models EffectorGeneP uses assembled transcripts mapped to the reference genome and from their exon/intron structure derives the longest ORF with its corresponding introns, 3’ and 5’ UTRs, Kozak sequence as well as intergenic sequences in the genome. To distinguish between CDSs and transcriptional noise, we also used the set of small genome-encoded ORFs (complete with both start and stop codon, ≤ 600 bps) as training data. Furthermore, we used the codon composition of these ORFs to generate a set of 10,000 random CDSs with lengths sampled from the lengths of the longest ORFs described above. The set of random CDS sequences is later also used to derive *p*-values in the gene annotation stage. We trained ML models to distinguish between the following types of genomic regions for subsequent gene annotation: the three types of CDSs, UTRs, intergenic regions, introns, small ORFs, random CDS sequences and Kozak sequences. For a given input sequence, EffectorGeneP derives 45 features as follows: GC content, proportion of third base (ACGT) in a codon, melting temperature (Biopython function Tm_NN), length, amino acid composition of the translation and local composition complexity^53^. Furthermore, EffectorGeneP calculates observed trinucleotide and hexanucleotide frequencies and derives the average observed/expected frequency for the sequence compared to CDS sequences, UTRs, introns and intergenic regions, respectively. From each of the three types of CDSs, EffectorGeneP also assesses codon usage by recording the frequencies of codons. For a given sequence, the deviation to the codon usage of CDSs encoding non-secreted proteins, CDSs encoding predicted secreted non-effector proteins and CDSs encoding predicted secreted effector proteins is calculated. We note that the presence of a stop codon is not part of the training set as a classifier could simply recognize the presence of one stop codon as a CDS. We used the WEKA tool box version 3.8.4^54^ to train an ensemble classifier composed of 67 logistic regression models and 67 C4.5 decision trees (J48 model in WEKA) (Supplementary Table 12). Soft voting is used to derive classifications and prediction probabilities for each classification. For the Kozak sequence ML models, we use an 11-dimensional feature vector that is composed of the GC content and each of the bases at the nine nucleotide positions before the start codon as well as the base after the start codon. Again, we trained an ensemble classifier composed of a logistic regression model and a C4.5 decision tree and used soft voting to derive prediction probabilities.

### Alignment and transcript calling for gene annotation

RNA-seq datasets were downloaded and reads were cleaned as described before. For gene annotation with EffectorGeneP, each replicate/sample was aligned separately with STAR 2.7.9a in 2-pass mode (-alignIntronMin 5 -- alignIntronMax 3000 --alignMatesGapMax 3000 --outFilterMultimapNmax 100) which removes junctions supported by <= two reads ^45^. Transcripts were assembled from the alignment with Stringtie 2.2.1 (-s 1 -m 200) with the appropriate strand setting ^46^ and merged into a consensus transcript set (stringtie –merge). To capture regions where transcripts were not called but that had RNA-seq evidence, we intersected this transcript set with the BAM file from the concatenated read mapping (bedtools intersect -v) ^55^ to extract mapped reads without an assembled transcript. We then called stringtie (-s 1 -m 200) for these mapped reads again to produce additional transcripts. Lastly, we concatenated the initial transcript set and the additional transcript set and converted this to a gff3 file with gffread ^56^ to input into EffectorGeneP for gene annotation.

### EffectorGeneP gene annotation from transcripts

First, each transcript is extended by 200 bp at both ends using the flanking reference genome sequence to compensate for incorrect transcript boundaries which we observed in some RNA-seq transcripts due to low coverage. Then, EffectorGeneP annotates genes from a given input transcript as follows: first, it extracts the longest ORF as well as smaller ORFs that are at least half its size. ORFs with the same stop codon but a different start site within 210 bps of the first start site are also kept if they are at least half of the size of the longest ORF. All these CDSs and their associated UTRs, introns (if present) and Kozak sequences are then classified by EffectorGeneP. For each potential CDS, the corresponding transcript segments receive a probability of constituting a CDS of a non-secreted protein, a CDS of a secreted protein, a CDS of an effector protein, an intron, intergenic sequence, a 3’ UTR, a 5’ UTR, a small ORF, a random CDS sequence or a Kozak sequence. From these probabilities, EffectorGeneP then derives the most likely gene model(s) per transcript as follows. First, EffectorGeneP only keeps those gene candidates with a CDS of > 0.5 probability, where the probability for a CDS is significant (*p* < 0.05). For the significance calculations, the 10,000 previously calculated random CDS sequences are used to derive *z*-scores and *p*-values from their CDS probabilities. Genes with introns that have average low probability (< 0.6) are dismissed. If a gene candidate passes these criteria, an overall gene score [0-100] is calculated using the probabilities of its CDS, introns (if present), UTRs and Kozak sequence (CDS weight 3; intron weight 6; 3’/5’ UTR weight 3; Kozak weight 3). EffectorGeneP also predicts signal peptides for all translated CDS sequences and assigns a higher score to those genes that encode proteins with a signal peptide and do not contain N-terminal transmembrane domains. By default, EffectorGeneP will use the transcript strand information as its most likely direction of transcription but it will also investigate gene candidates in the opposite direction of transcription. Single-exon gene candidates on the opposite strand are kept for further investigation, but if a multi-exon gene candidate has introns with average low probability (< 0.6) they will be dismissed. All gene candidates on the opposite strand receive a strand penalty in their final gene score to minimize false positive predictions. However, if a multi-exon gene candidate on the opposite strand has introns that contradict the strand direction it will be dismissed, as the strand of an intron can be reliably defined through the acceptor and donor sites. After collecting all gene candidates per transcript on both strands, a length-adjusted gene score is calculated for each one defined as the gene length (cds + intron lengths) multiplied by the gene score. Genes that encode secreted proteins will be assigned a higher score if they have a high effector coding probability. Secreted proteins are predicted as those that have a signal peptide (SignalP 4.1 -u 0.34 -U 0.34) and no transmembrane domains outside the N-terminal signal peptide region (TMHMM 2.0)^48,49^. EffectorGeneP can be run in default or conservative mode, where conservative mode implies more stringent thresholds for coding and gene scores. The gene with the highest length-adjusted gene score is returned as the final gene model for the transcript if its coding sequence and introns cover a minimum percentage of the transcript (default: 5%) and if the natural logarithm of the length-adjusted gene score is above a certain threshold (default: 5). If the UTRs of the highest-scoring gene are of sufficient length and they have a UTR probability of <= 0.98, one or both of them are investigated for transcript fusion events. Here, a more stringent gene filtering procedure is used where genes are only kept if the CDS probability is also higher than the intron and UTR probabilities (or alternatively if it has a high Kozak score > 0.5) and if the natural logarithm of the length-adjusted gene score is above a more stringent threshold (default: 5.7). Gene candidates are collected for the UTRs and each gene is represented by an interval comprised of its coding sequence start and coding sequence end, with 20 bps added on each side to provide space between adjacent genes. Each such interval has the length-adjusted gene score assigned as its weight. A dynamic programming algorithm is then used to find the maximum weight subset of genes such that no two genes overlap and these are returned as the final gene candidates for the transcript. Lastly, for each contig duplicate gene annotations as well as single-exon genes that are fully or partially contained in longer single-exon genes are removed.

### Benchmarking of gene annotation tools

We ran CodingQuarry 2.0 in pathogen mode^8^, Funannotate 1.8.5 (train with --jaccard_clip; predict with --optimize_augustus)^57^, Helixer with lineage fungi and default parameters^58^, TransDecoder 5.5.0 (https://github.com/TransDecoder/TransDecoder) and BRAKER3 (3.0.7) with parameters --fungus -- skip_fixing_broken_genes and the alignment file as input^59^. In addition, we obtained AUGUSTUS^60^, GeneMark^61^, PASA^62^, GlimmerHMM^63^ and SNAP^64^ predictions from the Funannotate intermediate files. De novo repeats were predicted with RepeatModeler 2.0.2a and the option -LTRStruct^65^. RepeatMasker 4.1.2p1 (-s -engine ncbi) (http://www.repeatmasker.org) was run with the RepeatModeler library to obtain statistics about repetitive element content. For gene annotation, RepeatMasker was run with the RepeatModeler library and the options -s (slow search) - nolow (does not mask low_complexity DNA or simple repeats) -engine ncbi. Secreted proteins were predicted as those that have a signal peptide (SignalP 4.1 -u 0.34 -U 0.34) and no transmembrane domains outside the N-terminal signal peptide region (TMHMM 2.0)^48,49^. Effectors were predicted with EffectorP 3.0^50^. For *B. graminis* f. sp. *tritici*, we extracted secreted proteins with an [YFW]xC or Y(x)xC sequence motif in their N-terminal region defined as the first 30 aas after the signal peptide cleavage site. For secretome comparisons to the reference annotations, we annotated secretomes with EffectorGeneP in conservative mode and removed genes that overlap with repetitive elements using bedtools coverage^55^ and a threshold of > 90% repeat coverage.

### Protein structure prediction and structural clustering

Structures of mature secreted proteins were predicted with Boltz-2 (2.2.0) and default parameters^24,25^, providing a multiple sequence alignment file for each protein generated by MMseqs2 (17-b804f) ^66^ against UniRef90 (downloaded on 23-09-2025) ^67^. Structural clustering was performed using FoldSeek v10 (-s 7.5 --tmalign-fast 0 --cov-mode 0 -c 0.5 --alignment-type 1 --tmscore-threshold 0.55) ^68^. The following PDB deposited resolved protein structures were included in the clustering: 2LW6, 2MM0, 2MYV, 2N37, 2N4O, 5A6W, 5ZLV, 6FMB, 6Q76, 7PP2, 7ZJY, 7ZK0, 7ZKD, 8C8A, 8OXH, 8OXI, 8OXJ, 8OXK, 8OXL.

### *AvrSr26* protoplast screen and validation

The two *P. graminis* f. sp. *tritici* library pools (part A and part B, containing 718 and 655 clones, respectively) described previously^27,28^ were screened in wheat protoplasts and differential expression analysis was conducted as described previously ^28^. Pooled libraries were propagated in ElectroMAX™ Stbl4™ competent cells and DNA was isolated using the QIAGEN EndoFree Plasmid Giga kit, as described previously ^27^. The reporter (pTA22-YFP), empty vector (pTA22), control (pTA22-Sr50) and pTA22-Sr26 plasmids were isolated from E. coli DH5α using the Macherey Nagel NucleoBond Xtra Maxi Plus EF kit. A total of 200,000 protoplast cells of wheat cultivar Fielder were transformed with a pooled effector library at a multiplicity of transfection (MOT) of 0.14 million molecules per cell for each construct along with either the empty vector pTA22, pTA22-Sr50 or pTA22-Sr26 (MOT = 36 million molecules per cell, 36, 57 and 82 µg, respectively) in triplicate as described^27^. A control protoplast sample (50,000 cells) was transformed with pTA22-YFP (10 µg; MOT = 36 million molecules per cell) alone to assess transfection efficiency by flow cytometry. Messenger (m)RNA was extracted from transformed protoplasts after 24 hours using the NEB Magnetic mRNA Isolation Kit (#S1550S). Library-specific cDNA synthesis and PCR was carried out on 30 ng mRNA using the Invitrogen SuperScript IV One-Step RT-PCR System with ezDNase kit (Cat. # 12595100) with the forward and reverse primers ZmUbi1_5UTR_F3b (5’GCACACACACACAACCAG-3’) and FS_cDNA_R (5’- TGCTAGATCTCGACAGTACG-3’) as previously described^27^. Illumina libraries were generated for each cDNA sample (three replicates of each treatment and control) using the Illumina DNA Prep Kit (Cat. # 20060059) and IDT for Illumina DNA/RNA UD Indexes Set A (Cat. # 20027213) following the manufacturer’s protocol. Illumina libraries were sequenced using the NextSeq2000 at the ACRF Biomolecular Resource Facility, The John Curtin School of Medical Research, Australian National University using the NextSeq P1 300 cycles kit, with 150 bp paired end reads. Sequencing reads were cleaned using fastp v0.23.2 and default parameters. The clean reads were aligned to the coding sequences of the effector candidates with STAR76 2.7.9 (--alignIntronMax 1, --outFilterMultimapNmax 10) ^45^. Read counts were compiled with samtools idxstats^69^ and imported into DESeq for differential expression analysis using default parameters followed by lfcShrink (type=“apeglm”) to compare the resistance gene and empty vector treatments^70^. Volcano plots were produced with EnhancedVolcano (https://github.com/kevinblighe/EnhancedVolcano).

For the alleles of the positive clones, the coding sequences of the mature proteins were codon optimized for wheat and synthesised directly in pDONR207 (Epoch Life Science) and cloned into the expression vector pTA22 ^27^ via a Gateway LR reaction (Invitrogen). For transient expression of *Sr26* and *AvrSr26* variants in wheat protoplasts, plasmids were extracted using the NucleoBond Xtra Midi EF kit (Macherey-Nagel). Wheat seedlings (cv Fielder) were grown in Martins Seed Raising and Cutting Mix at 24°C under a 12 h light (∼130 µmol m-2 s-1) and 12 h dark cycle. At seven days, the first leaf was harvested for protoplast isolation and cells transformed as previously described with minor modifications ^27^. Transfected cells were incubated in 2ml Eppendorf tubes for 20h. Following the incubation, half of the volume of supernatant was removed, and the protoplasts resuspended in the remainder. Fluorescence was then measured with a CLARIOstar Plus plate reader (BMG Labtech; the top optic was used, optic settings: presetname = YFP, excitation = 497-15, dichroic filter = 517.2, emission = 540-20, focal height = 5.7mm, scan settings: number of flashes per scan = 4, scan mode = matrix scan, scan matrix dimension = 30x30, scan width = 7mm). The gain was set based on fluorescence scanning of YFP control samples (scan settings as above except: number of flashes per scan = 100, scan mode = spiral average). Fluorescence was normalized against the mean of the YFP control samples.

### AvrSr26 transient expression in N. benthamiana and N. tabacum

*N. benthamiana* and *N. tabacum* plants were grown at 24°C with 16 h light. Four-week-old plants were used for transient expression assays. *Sr26* was cloned into pAM-PAT-GW-YFPv vector with a YFP tag fused at the C terminus. *AvrSr26* candidates were cloned into pBIN19-YFP-GW vector with a YFP tag fused at the N-terminus. *Agrobacterium* strains GV3103 pMP90 RK (pAM-Sr26-YFPv) and GV3101 pMP90 RK (pBIN19-YFP-AvrSr26) were cultured overnight at 28°C in LB media with required antibiotic selections. Cells were harvested and resuspended in buffer (10 mM MES pH 5.6, 10 mM MgCl_2_, 150 μM acetosyringone) and incubated at room temperature for 2 h. For HR assay, the Sr26-containing *Agrobacterium* were infiltrated at optical density (OD_600_) of 0.3, and the AvrSr26-containing *Agrobacterium* were infiltrated at OD_600_ = 0.5. All leaves were documented 3 days after infiltration and cell death on *N. benthamiana* leaves were scored visually on a 0-5 scale.

### Virus-mediated *in planta AvrSr26* effector variants expression assay

The barley stripe mosaic virus (BSMV) mediated protein overexpression system (BSMV-VOX), comprising three T-DNA binary plasmids, pCaBS-α, pCaBS-β, and pCa-γb2A-LIC, was used ^71^. Mature coding sequences of the *AvrSr26* variants (#0998, #1000, #1007, and #1033), in which the predicted signal peptides were replaced by a methionine start codon, were PCR-amplified using primers flanking each coding sequence and carrying 5′ extensions (forward primer: CCAACCCAGGACCGTTGATG; reverse primer: AACCACCACCACCGTTA) to enable ligation-independent cloning into pCa-γb2A-LIC (Supplementary Table 13). Amplified fragments were cloned into pCa-γb2A-LIC following the procedure of Kanyuka (2022), and all constructs were verified by Sanger sequencing. The BSMV binary plasmids were transformed into *Agrobacterium tumefaciens* strain GV3101(pMP90). Agroinfiltration of 3-4-week-old *Nicotiana benthamiana* plants was performed as described previously^71^. Crude sap from infiltrated *N. benthamiana* leaves was subsequently used to mechanically inoculate leaves of three-leaf-stage wheat plants of cultivars Avocet (*Sr26*) and Cadenza (*sr26*) ^72^. BSMV:iLov served as a negative control ^73^. Inoculated wheat plants were maintained in a controlled-environment growth room at 23°C/20°C day/night temperatures, 60-65% relative humidity, and a 16-h photoperiod with ∼220 μmol m^-2^ s^-1^ light intensity. The presence or absence of viral symptoms in systemic leaves was assessed at 10, 16, and 24 days post-inoculation. For each time point, the number of plants displaying systemic symptoms was recorded relative to the total number inoculated. Each experiment included 8-10 individual plants per BSMV-VOX construct, with two replicates of the experiment performed.

### *AvrSr26-*like gene family analysis

Sequence homologs were extracted by searching the four positive library clones against proteomes with phmmer (-E 0.001 --max)^74^. Mature secreted proteins were aligned with muscle 5.0.1428^75^ and visualized in Jalview^76^. Alignment identity was extracted with esl-alistat^74^. A phylogenetic tree was constructed with fasttree 2.1.11^77^ from the protein alignment and visualized in iTol ^78^.

### Stem rust RNA-seq and PacBio Iso-Seq data generation

Morocco/Fielder seedlings were grown in two pots with five plants per pot and infected with *P. graminis* f. sp. *tritici* 21-0 at 10 days after sowing. For each plant the first leaf was collected at 6 days post infection and frozen in liquid nitrogen. RNA was extracted with the Maxwell® RSC Plant RNA Kit (Promega, Catalog #AS1500) and the Maxwell robot. RNA and Iso-Seq data was generated from this material by AGRF (Brisbane, Australia). We processed the Iso-Seq reads with isoseq refine to remove polyA tails and artificial concatemers (https://github.com/PacificBiosciences/pbbioconda). We then cleaned these reads using fastp (--trim_poly_x) and an adapter file containing Iso-Seq primers ^44^. Iso-Seq reads were aligned to the genome with minimap2 (-ax splice:hq -uf -G 3000)^79^ and long-reads transcripts were called with Stringtie 2.2.1 (-L -m 200)^46^.

The RNA-seq reads were aligned with STAR 2.7.9a in 2-pass mode (-alignIntronMin 5 --alignIntronMax 3000 -- alignMatesGapMax 3000 --outFilterMultimapNmax 100) which removes junctions supported by <= two reads^45^. RNA-seq transcripts were assembled from the alignment with Stringtie 2.2.1 (-s 1.5 -m 200 --rf)^46^ and hybrid RNA-seq and Iso-Seq transcript were assembled from the alignment with Stringtie 2.2.1 (-s 1.5 -m 200 --mix). Transcripts that overlap with repetitive elements where removed using bedtools coverage^55^ and a threshold of > 90% repeat coverage.

### Stem rust PacBio genome sequencing and assembly

High molecular DNA from *P. graminis* f. sp. *tritici* isolate Pgt21-0 urediniospores was extracted as previously described ^26,80^. DNA quality was assessed with a NanoDrop spectrophotometer (Thermo Scientific) and the concentration quantified using a broad-range assay in a Qubit 3.0 fluorometer (Invitrogen). DNA library preparation (10-to 15-kbp fragments; Pippin Prep) and sequencing on the PacBio Sequel II platform (One SMRT Cell 8M) were performed by the Australian Genome Research Facility (St Lucia, Queensland, Australia) following manufacturer guidelines. Hi-C reads were previously generated^26^. The HiFi reads were assembled using hifiasm 0.19.6 in Hi-C integration mode and with default parameters^81^. Contaminants were identified using sequence similarity searches (BLAST 2.11.0 -db nt -evalue 1e-5 -perc_identity 75)^82^. HiFi reads were aligned to the assembly with minimap2 v.2.22 (-ax map-hifi -secondary=no)^79^, and contig coverage was called using bbmap’s pileup.sh tool on the minimap2 alignment file (http://sourceforge.net/projects/bbmap). All contaminant contigs, contigs with less than 15× coverage, and the mitochondrial contigs were removed from the assembly. Chromosomes were curated using visual inspection of Hi-C contact maps produced using Hi-C-Pro 3.1.0 (MAPQ = 10)^83^ and HiCExplorer 3.7.2^84^. An incorrect inversion on chromosome 3A in the previously generated PacBio RSII assembly^26^ was detected and is corrected in this assembly.

## Data availability

The EffectorGeneP software is available at https://github.com/JanaSperschneider/EffectorGeneP and model files are available at https://effectorp.csiro.au/effectorgenep.html. Sequencing data (PacBio HiFi data, 6 dpi RNA-seq and Iso-Seq) and the PacBio HiFi genome assembly as well as the PacBio RSII assembly with the corrected chromosome 3A for Pgt21-0 is available at the CSIRO Data Access Portal: https://data.csiro.au/collection/csiro:72194. Protoplast library screening data is available at the CSIRO Data Access Portal: https://data.csiro.au/collection/csiro:72195.

## Contributions

J.S. conceptualized the study, curated input data, designed the methodology and machine learning models, performed computational experiments and implemented the software. C.L.P., J.C., J.L. and D.L. performed effector validation experiments. T.A. and C.B. performed protoplast library screening experiments. E.C.H. and D.L. performed transcriptional data generation. D.L. performed PacBio genome sequencing data generation. J.S., K.K., T.V., M.F. and P.N.D supervised the work, managed activities and acquired funding. J.S. wrote the original draft and all authors reviewed and edited the manuscript.

## References

1. Freedman, A. H. & Sackton, T. B. Building better genome annotations across the tree of life. Genome Res. 35, 1261–1276 (2025).

2. Stiehler, F. et al. Helixer: cross-species gene annotation of large eukaryotic genomes using deep learning. Bioinformatics 36, 5291–5298 (2021).

3. Holst, F. et al. Helixer–<em>de novo</em> Prediction of Primary Eukaryotic Gene Models Combining Deep Learning and a Hidden Markov Model. bioRxiv 2023.02.06.527280 (2023) doi:10.1101/2023.02.06.527280.

4. Gabriel, L., Becker, F., Hoff, K. J. & Stanke, M. Tiberius: End-to-End Deep Learning with an HMM for Gene Prediction. bioRxiv 2024.07.21.604459 (2024) doi:10.1101/2024.07.21.604459.

5. Scalzitti, N., Jeannin-Girardon, A., Collet, P., Poch, O. & Thompson, J. D. A benchmark study of ab initio gene prediction methods in diverse eukaryotic organisms. BMC Genomics 21, 293 (2020).

6. Gabriel, L. et al. BRAKER3: Fully automated genome annotation using RNA-seq and protein evidence with GeneMark-ETP, AUGUSTUS and TSEBRA. bioRxiv 2023.06.10.544449 (2024) doi:10.1101/2023.06.10.544449.

7. Tang, S., Lomsadze, A. & Borodovsky, M. Identification of protein coding regions in RNA transcripts. Nucleic Acids Res 43, e78 (2015).

8. Testa, A. C., Hane, J. K., Ellwood, S. R. & Oliver, R. P. CodingQuarry: highly accurate hidden Markov model gene prediction in fungal genomes using RNA-seq transcripts. BMC genomics 16, 170 (2015).

9. Arendsee, Z. W., Li, L. & Wurtele, E. S. Coming of age: orphan genes in plants. Trends in Plant Science 19, 698–708 (2014).

10. Lo Presti, L. et al. Fungal effectors and plant susceptibility. Annual review of plant biology 66, 513–45 (2015).

11. Sperschneider, J. et al. Advances and challenges in computational prediction of effectors from plant pathogenic fungi. PLoS pathogens 11, e1004806 (2015).

12. Testa, A. C. Effector gene prediction from fungal pathogen genome assemblies. in (2016).

13. Rehmany, A. P. et al. Differential Recognition of Highly Divergent Downy Mildew Avirulence Gene Alleles by *RPP1* Resistance Genes from Two Arabidopsis Lines. Plant Cell 17, 1839–1850 (2005).

14. Torto, T. A. et al. EST Mining and Functional Expression Assays Identify Extracellular Effector Proteins From the Plant Pathogen Phytophthora. Genome Res. 13, 1675–1685 (2003).

15. Haas, B. J. et al. Genome sequence and analysis of the Irish potato famine pathogen Phytophthora infestans. Nature 461, 393–8 (2009).

16. Stam, R. et al. Identification and Characterisation CRN Effectors in Phytophthora capsici Shows Modularity and Functional Diversity. PLoS ONE 8, e59517 (2013).

17. Tabima, J. F. & Grünwald, N. J. *effectR* : An Expandable R Package to Predict Candidate RxLR and CRN Effectors in Oomycetes Using Motif Searches. MPMI 32, 1067–1076 (2019).

18. Lapalu, N. et al. Improved gene annotation of the fungal wheat pathogen <em>Zymoseptoria tritici</em> based on combined Iso-Seq and RNA-Seq evidence. bioRxiv 2023.04.26.537486 (2023) doi:10.1101/2023.04.26.537486.

19. Sperschneider, J. et al. Nuclear exchange generates population diversity in the wheat leaf rust pathogen Puccinia triticina. Nat Microbiol 8, 2130–2141 (2023).

20. Saitoh, H. et al. Large-Scale Gene Disruption in Magnaporthe oryzae Identifies MC69, a Secreted Protein Required for Infection by Monocot and Dicot Fungal Pathogens. PLoS pathogens 8, e1002711 (2012).

21. Khrunyk, Y., Münch, K., Schipper, K., Lupas, A. N. & Kahmann, R. The use of FLP-mediated recombination for the functional analysis of an effector gene family in the biotrophic smut fungus Ustilago maydis. New Phytologist 187, 957–968 (2010).

22. Tucker, S. L. et al. A Fungal Metallothionein Is Required for Pathogenicity of *Magnaporthe grisea*. Plant Cell 16, 1575–1588 (2004).

23. Syme, R. A. et al. Comprehensive Annotation of the Parastagonospora nodorum Reference Genome Using Next-Generation Genomics, Transcriptomics and Proteogenomics. PLoS One 11, e0147221 (2016).

24. Wohlwend, J. et al. Boltz-1 Democratizing Biomolecular Interaction Modeling. Preprint at 10.1101/2024.11.19.624167 (2024).

25. Passaro, S. et al. Boltz-2: Towards Accurate and Efficient Binding Affinity Prediction. Preprint at 10.1101/2025.06.14.659707 (2025).

26. Li, F. et al. Emergence of the Ug99 lineage of the wheat stem rust pathogen through somatic hybridisation. Nat Commun 10, 5068 (2019).

27. Arndell, T. et al. Pooled effector library screening in protoplasts rapidly identifies novel Avr genes. Nat. Plants 10, 572–580 (2024).

28. Chen, R. et al. A wheat tandem kinase activates an NLR to trigger immunity. Science 387, 1402–1408 (2025).

29. Shumate, A., Wong, B., Pertea, G. & Pertea, M. Improved transcriptome assembly using a hybrid of long and short reads with StringTie. PLoS Comput Biol 18, e1009730 (2022).

30. Upadhyaya, N. M. et al. Genomics accelerated isolation of a new stem rust avirulence gene–wheat resistance gene pair. Nat. Plants 7, 1220–1228 (2021).

31. Haas, B. J. et al. Automated eukaryotic gene structure annotation using EVidenceModeler and the Program to Assemble Spliced Alignments. Genome Biol 9, R7 (2008).

32. Li, G., Dulal, N., Gong, Z. & Wilson, R. A. Unconventional secretion of Magnaporthe oryzae effectors in rice cells is regulated by tRNA modification and codon usage control. Nat Microbiol 8, 1706–1716 (2023).

33. Davison, H. R. et al. The promise of AlphaFold for gene structure annotation. Preprint at 10.1101/2025.10.21.683479 (2025).

34. Marín, M., Uversky, V. N. & Ott, T. Intrinsic Disorder in Pathogen Effectors: Protein Flexibility as an Evolutionary Hallmark in a Molecular Arms Race. The Plant Cell 25, 3153–3157 (2013).

35. Shen, D., Li, Q., Sun, P., Zhang, M. & Dou, D. Intrinsic disorder is a common structural characteristic of RxLR effectors in oomycete pathogens. Fungal Biology 121, 911–919 (2017).

36. Zhang, J. et al. A recombined Sr26 and Sr61 disease resistance gene stack in wheat encodes unrelated NLR genes. Nat Commun 12, 3378 (2021).

37. Periyannan, S., Milne, R. J., Figueroa, M., Lagudah, E. S. & Dodds, P. N. An overview of genetic rust resistance: From broad to specific mechanisms. PLoS Pathog 13, e1006380 (2017).

38. Zhang, J., Zhang, P., Karaoglu, H. & Park, R. F. Molecular Characterization of Australian Isolates of *Puccinia graminis* f. sp. *tritici* Supports Long-Term Clonality but also Reveals Cryptic Genetic Variation. Phytopathology® 107, 1032–1038 (2017).

39. Mago, R. et al. Transfer of stem rust resistance gene SrB from Thinopyrum ponticum into wheat and development of a closely linked PCR-based marker. Theor Appl Genet 132, 371–382 (2019).

40. Kunz, L. et al. Dual recognition of structurally unrelated mildew effectors underlies the broad-spectrum resistance of Pm3e in wheat. Preprint at 10.1101/2025.10.26.683672 (2025).

41. Redkar, A. et al. The Arabidopsis *WRR4A* and *WRR4B* paralogous NLR proteins both confer recognition of multiple *Albugo candida* effectors. New Phytologist 237, 532–547 (2023).

42. Zhao, H. et al. Structural basis for the broad recognition specificity of an Arabidopsis immune receptor. Preprint at 10.64898/2026.02.04.703898 (2026).

43. Silverstein, K. A. T. et al. Small cysteine-rich peptides resembling antimicrobial peptides have been under-predicted in plants. The Plant Journal 51, 262–280 (2007).

44. Chen, S., Zhou, Y., Chen, Y. & Gu, J. fastp: an ultra-fast all-in-one FASTQ preprocessor. Bioinformatics 34, i884–i890 (2018).

45. Dobin, A. et al. STAR: ultrafast universal RNA-seq aligner. Bioinformatics 29, 15–21 (2013).

46. Pertea, M. et al. StringTie enables improved reconstruction of a transcriptome from RNA-seq reads. Nat Biotechnol 33, 290–295 (2015).

47. Fu, L., Niu, B., Zhu, Z., Wu, S. & Li, W. CD-HIT: accelerated for clustering the next-generation sequencing data. Bioinformatics 28, 3150–3152 (2012).

48. Krogh, A., Larsson, B., von Heijne, G. & Sonnhammer, E. L. Predicting transmembrane protein topology with a hidden Markov model: application to complete genomes. Journal of molecular biology 305, 567–80 (2001).

49. Petersen, T. N., Brunak, S., von Heijne, G. & Nielsen, H. SignalP 4.0: discriminating signal peptides from transmembrane regions. Nature methods 8, 785–6 (2011).

50. Sperschneider, J. & Dodds, P. N. EffectorP 3.0: Prediction of Apoplastic and Cytoplasmic Effectors in Fungi and Oomycetes. MPMI 35, 146–156 (2022).

51. Singh, U. & Wurtele, E. S. orfipy: a fast and flexible tool for extracting ORFs. Bioinformatics 37, 3019–3020 (2021).

52. Rice, P., Longden, I. & Bleasby, A. EMBOSS: the European Molecular Biology Open Software Suite. Trends in genetics : TIG 16, 276–7 (2000).

53. Konopka, A. K. Sequence Complexity and Composition. in eLS (2005). doi:10.1038/npg.els.0005260.

54. Mark Hall et al. The WEKA data mining software: an update. SIGKDD Explor. Newsl. 11, 10–18 (2009).

55. Quinlan, A. R. & Hall, I. M. BEDTools: a flexible suite of utilities for comparing genomic features. Bioinformatics 26, 841–842 (2010).

56. Pertea, G. & Pertea, M. GFF Utilities: GffRead and GffCompare. F1000Res 9, ISCB Comm J–304 (2020).

57. Palmer, J. M. & Stajich, J. Funannotate v1.8.1: Eukaryotic genome annotation. Zenodo (2020).

58. Holst, F. et al. Helixer– de novo Prediction of Primary Eukaryotic Gene Models Combining Deep Learning and a Hidden Markov Model. Preprint at 10.1101/2023.02.06.527280 (2023).

59. Gabriel, L. et al. BRAKER3: Fully automated genome annotation using RNA-seq and protein evidence with GeneMark-ETP, AUGUSTUS, and TSEBRA. Genome Res. 34, 769–777 (2024).

60. Stanke, M., Schöffmann, O., Morgenstern, B. & Waack, S. Gene prediction in eukaryotes with a generalized hidden Markov model that uses hints from external sources. BMC Bioinformatics 7, 62 (2006).

61. Ter-Hovhannisyan, V., Lomsadze, A., Chernoff, Y. O. & Borodovsky, M. Gene prediction in novel fungal genomes using an ab initio algorithm with unsupervised training. Genome Res. 18, 1979–1990 (2008).

62. Haas, B. J. Improving the Arabidopsis genome annotation using maximal transcript alignment assemblies. Nucleic Acids Research 31, 5654–5666 (2003).

63. Majoros, W. H., Pertea, M. & Salzberg, S. L. TigrScan and GlimmerHMM: two open source ab initio eukaryotic gene-finders. Bioinformatics 20, 2878–2879 (2004).

64. Korf, I. Gene finding in novel genomes. BMC Bioinformatics 5, 59 (2004).

65. Flynn, J. M. et al. RepeatModeler2 for automated genomic discovery of transposable element families. Proc Natl Acad Sci USA 117, 9451–9457 (2020).

66. Steinegger, M. & Söding, J. MMseqs2 enables sensitive protein sequence searching for the analysis of massive data sets. Nat Biotechnol 35, 1026–1028 (2017).

67. The UniProt Consortium et al. UniProt: the Universal Protein Knowledgebase in 2025. Nucleic Acids Research 53, D609–D617 (2025).

68. Van Kempen, M. et al. Fast and accurate protein structure search with Foldseek. Nat Biotechnol 42, 243–246 (2024).

69. Li, H. et al. The Sequence Alignment/Map format and SAMtools. Bioinformatics 25, 2078–2079 (2009).

70. Love, M. I., Huber, W. & Anders, S. Moderated estimation of fold change and dispersion for RNA-seq data with DESeq2. Genome biology 15, 550 (2014).

71. Kanyuka, K. Virus-Mediated Protein Overexpression (VOX) in Monocots to Identify and Functionally Characterize Fungal Effectors. in Effector-Triggered Immunity (eds Kufer, T. A. & Kaparakis-Liaskos, M.) vol. 2523 93–112 (Springer US, New York, NY, 2022).

72. Yu, L.-X. et al. Haplotype diversity of stem rust resistance loci in uncharacterized wheat lines. Mol Breeding 26, 667–680 (2010).

73. Franco-Orozco, B. et al. A new proteinaceous pathogen-associated molecular pattern (PAMP) identified in Ascomycete fungi induces cell death in Solanaceae. New Phytologist 214, 1657–1672 (2017).

74. Finn, R. D., Clements, J. & Eddy, S. R. HMMER web server: interactive sequence similarity searching. Nucleic acids research 39, W29–37 (2011).

75. Edgar, R. C. MUSCLE: multiple sequence alignment with high accuracy and high throughput. Nucleic acids research 32, 1792–7 (2004).

76. Waterhouse, A. M., Procter, J. B., Martin, D. M., Clamp, M. & Barton, G. J. Jalview Version 2--a multiple sequence alignment editor and analysis workbench. Bioinformatics 25, 1189–91 (2009).

77. Price, M. N., Dehal, P. S. & Arkin, A. P. FastTree 2 – Approximately Maximum-Likelihood Trees for Large Alignments. PLoS ONE 5, e9490 (2010).

78. Letunic, I. & Bork, P. Interactive Tree Of Life (iTOL): an online tool for phylogenetic tree display and annotation. Bioinformatics 23, 127–128 (2007).

79. Li, H. Minimap2: pairwise alignment for nucleotide sequences. Bioinformatics 34, 3094–3100 (2018).

80. Schwessinger, B. & Rathjen, J. P. Extraction of High Molecular Weight DNA from Fungal Rust Spores for Long Read Sequencing. Methods Mol Biol 1659, 49–57 (2017).

81. Cheng, H., Concepcion, G. T., Feng, X., Zhang, H. & Li, H. Haplotype-resolved de novo assembly using phased assembly graphs with hifiasm. Nat Methods 18, 170–175 (2021).

82. Altschul, S. F., Gish, W., Miller, W., Myers, E. W. & Lipman, D. J. Basic local alignment search tool. Journal of molecular biology 215, 403–10 (1990).

83. Servant, N. et al. HiC-Pro: an optimized and flexible pipeline for Hi-C data processing. Genome Biol 16, 259 (2015).

84. Ramírez, F. et al. High-resolution TADs reveal DNA sequences underlying genome organization in flies. Nat Commun 9, 189 (2018).

